# Dynamics of infant white matter maturation from birth to 6 months

**DOI:** 10.1101/2025.02.13.638114

**Authors:** Benjamin Risk, Longchuan Li, Warren Jones, Sarah Shultz

**Author notes:** Correspondence: Sarah Shultz, PhD, Associate Professor Department of Pediatrics, Emory University School of Medicine, Marcus Autism Center, Children’s Healthcare of Atlanta /.

## Abstract

The first months after a baby’s birth encompass the most rapid period of postnatal change in the human lifespan, but longitudinal trajectories of white matter maturation in this period remain uncharted. Using densely sampled diffusion tensor images collected longitudinally at a mean rate of 1 scan per 1.55 days, we measured non-linear growth and growth rate trajectories of major white matter tracts from birth to 6 months. Growth rates at birth were 6 to 11 times faster than at 6 months, with tracts less mature at birth developing fastest. When matched on chronological age, shorter gestation infants had less mature white matter at birth but faster growth rates than their longer gestation peers; however, growth trajectories were highly similar when corrected for gestational age. This is the first study to estimate white matter trajectories using dense sampling in the first 6 post-natal months, which can inform the study of neurodevelopmental disorders beginning in infancy.

## Introduction

The development of white matter tracts—the bundles of axons that carry information between neurons—is a complex and dynamic process that begins in the fetal period and continues through adulthood^1^, with the largest changes occurring during the first year of life^2,3^. During the first postnatal year, white matter tracts undergo rapid myelination, which enables fast transmission of electrical signals between neural systems^4,5^ and establishes the structural foundation for functionally organized brain networks^6^. This dynamic change is accompanied by extremely rapid cerebral growth and maturation, with total brain volume doubling^7^ and synaptic density quadrupling within year one alone^8^. Despite the importance of the first months of postnatal development, temporally precise growth and growth-rate trajectories of white matter development during this period are uncharted.

The extraordinary rate of growth in these first postnatal months contributes to this being a sensitive period of development, when neural circuitry is especially susceptible to endogenous and exogenous influences^9–13^. This concept of enhanced susceptibility during periods of rapid brain growth originated from decades of animal research demonstrating that brain regions that are growing most rapidly are also especially vulnerable to influences from a wide range of factors including but not limited to stress, nutrition, sensory deprivation, and hormonal changes^14–18^. Given that white matter tracts mature asynchronously—with some tracts maturing earlier and/or more quickly than others^19^—the vulnerability of those tracts may also be highly variable, with different tracts and their associated behaviors being more or less vulnerable at different times in early development. A consequence of that variability is that quantification of longitudinal, temporally precise growth and growth-rate trajectories of white matter development can provide a critical resource for future studies identifying tract-specific periods of both heightened vulnerability (in the face of stressors or insults) or heightened opportunity (when adaptive influences may be of greatest benefit).

A key factor that may impact white matter growth and growth-rate trajectories is gestational age at birth. Shortened gestation, even amongst term-born infants, reduces white matter maturity^22,23^ and is associated with increased likelihood for neurodevelopmental disability^24,25^. However, the impact throughout infants’ first 6 postnatal months is largely unknown. Three hypotheses have been proposed: 1) that early exposure to the extrauterine environment accelerates white matter maturation^26,27^; 2) that growth-rates of white matter development are largely unaltered by early exposure to the extrauterine environment^28,29^; and 3) that birth itself may slow trajectories of white matter development, potentially leading to a persistent reduction in white matter maturity amongst infants with shorter gestation.^30^

Here we show that the dynamics of white matter maturation are time-varying, asynchronous, and impacted by infant gestational age at birth. Growth rates of diffusion tensor imaging (DTI) metrics peak at birth and slow thereafter, with less mature tracts at birth developing more rapidly over the first 6 months. Earlier gestational age at birth was associated with reduced maturation. We show that infants born at shorter gestation had faster growth rates when matched on chronological age, but similar growth rates when corrected for gestational age at birth. This supports the unaltered pace hypothesis.

These longitudinal profiles offer new insight into the complex and dynamic processes underlying white matter development in the human brain and provide temporally precise benchmarks against which to measure how departures from typical trajectories can lead to disability.

## Results

In the present study, we used a very densely sampled longitudinal design to study white matter tract development in infants’ first 6 postnatal months. Diffusion MRI data were collected using a non-uniform longitudinal sampling design, with data collected from each infant at up to 3 randomized time points between birth and 6 months from 79 largely term-born typically developing infants (87% of sample ≥ 37 weeks’ gestation, see Supplementary Table 1 for participant demographics). This approach yielded dense coverage over the 0-to 6-month period (129 scans, each separated by a mean of only 1.55 days (stdev=1.69, min=0 days, max=9 days), a necessity for generating temporally precise trajectories during periods of rapid developmental change^20^. See Supplementary Fig.1 for a distribution of participant age at each scan. Eleven major white matter tracts were identified using whole-brain tractography. Generalized additive mixed models (GAMMs), an approach that models response variables as smooth functions of predictors, without a priori trajectory shape assumptions, were used to model growth and change rate trajectories of four DTI-derived metrics: fractional anisotropy (FA), mean diffusivity (MD), axial diffusivity (AD), and radial diffusivity (RD).

### Whole-Brain Growth Trajectories and Growth Rates

The first 6 months of life were characterized by increasing FA and decreasing diffusivity measures (MD, AD, RD) (Fig. 1). For all DTI metrics, the maximum growth rates occurred at birth, with FA increasing by approximately 0.44% per day, and MD, AD, and RD decreasing by-0.34%,-0.25% and-0.39%, respectively. In FA, the rate of change decreases quickly until approximately 100 days, at which point it is 0.11%/day, and is relatively stable from 150 to 200 days increasing at just over 0.05%/day. In MD, AD, and RD, the negative growth rate rapidly slows until roughly 70 days, at which point the growth rate is-0.10%,-0.08% and-0.12%, respectively, then levels off, with growth rates around-0.04%,-0.03% and-0.05% from 160 to 200 days. Growth rate is nearly six times faster at birth than at 6 months in FA and approximately 11 times faster at birth than at 6 months in MD, AD, and RD (Supplementary Fig. 4). Thus, the dynamics of white matter development change substantially within the first 6 postnatal months.

**Figure 1.**
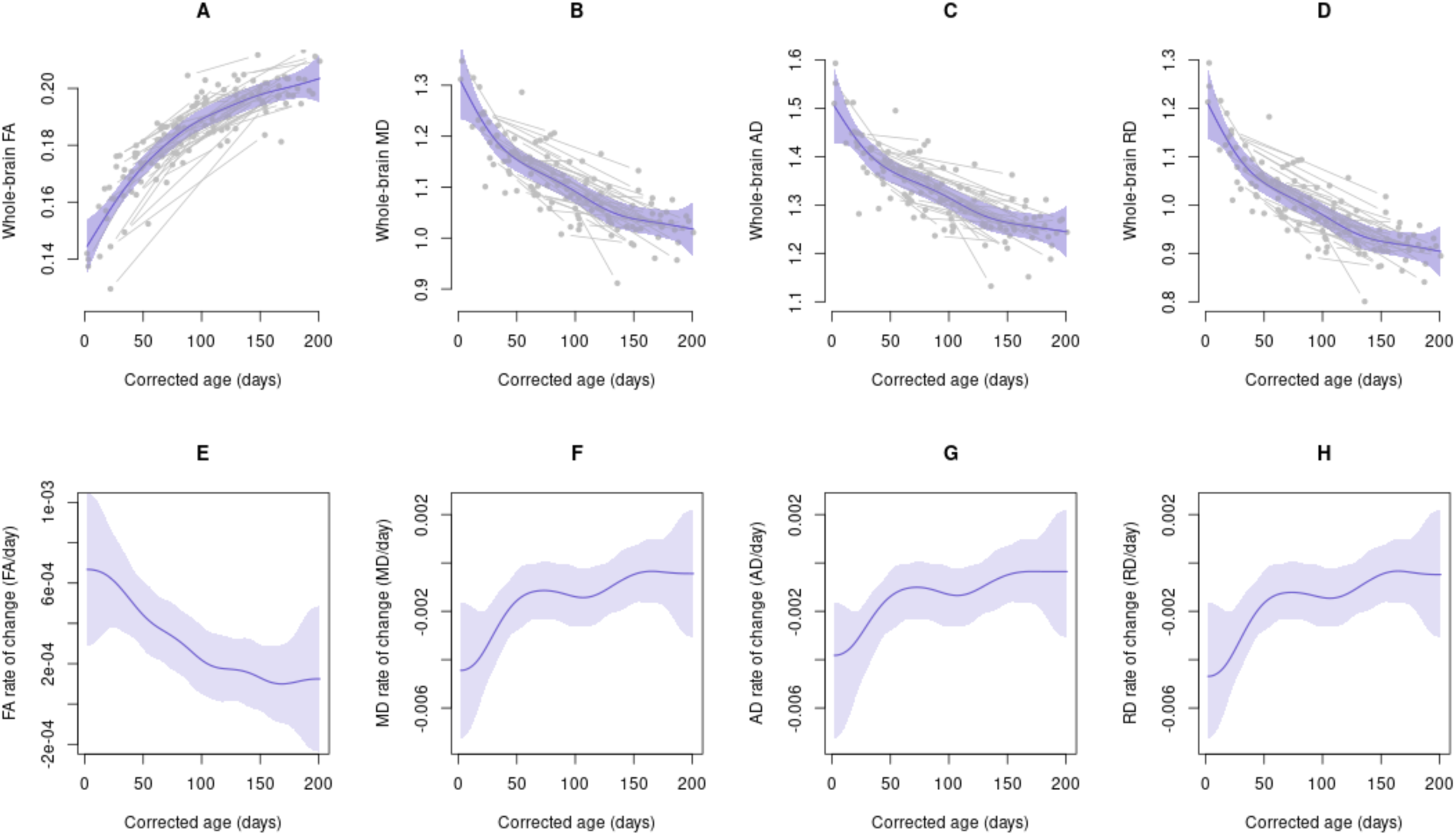
**Growth curves (A-D) and growth rates (E-H) in whole-brain FA, MD, AD, and RD in the first 6 postnatal months**. Gray points represent the raw data with lines connecting points from the same infant. Blue lines represent the fitted mean curves **(A-D)** and their first derivatives **(E-H)**. Bands represent the 95% simultaneous confidence bands.

### Regional Growth Trajectories and Growth Rates

Similar to growth patterns of whole-brain white matter, all white matter pathways had increasing FA and decreasing diffusivity measures over the first 6 postnatal months (Fig. 2). Most tracts had their maximum growth rates at birth with lower growth rates at later mo e Supplementary Fig. 5 for growth curves and growth rates plotted with 95% simultaneous confidence bands). Some tracts exhibited large changes in growth rates over the first six months (e.g., the body of the corpus callosum, (CCb)), while a few tracts exhibited smaller dynamics (e.g., nearly constant growth rates for the fornix (Fx)). For most tracts, the growth rate at 100 days was less than half the growth rate at birth, and the rates began to stabilize around month 5.

**Figure 2.**
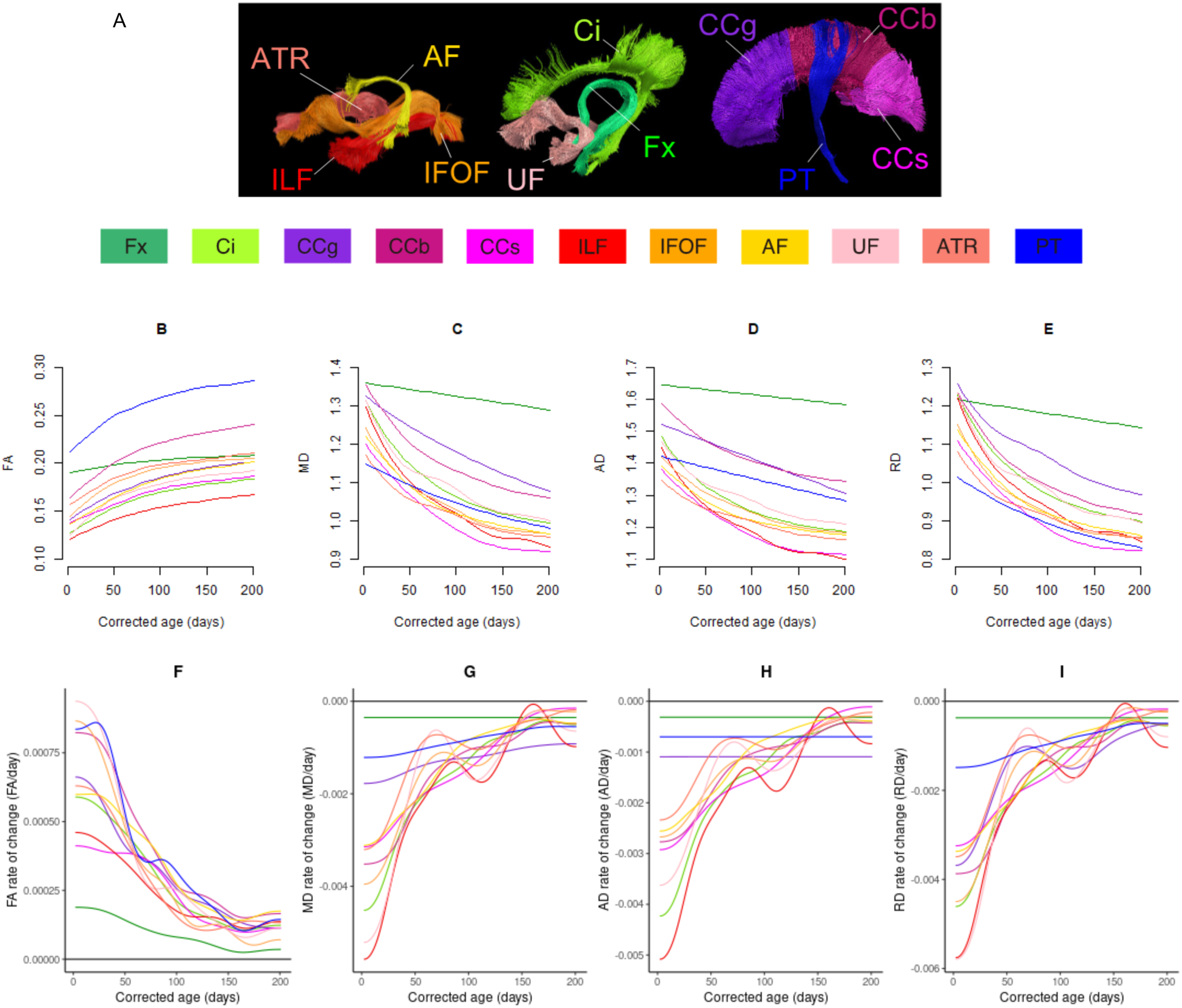
**Growth curves and growth rates of FA, MD, AD, and RD in 11 white matter pathways during the first 6 postnatal months**. **(A)** Delineation of 11 major white matter pathways from whole-brain tractography. **AF**: arcuate fasciculus; **ATR**: anterior thalamic radiation; **CCb**: body of corpus callosum; **CCg**: genu of corpus callosum; **CCs**: splenium of corpus callosum; **Ci**: cingulum; **Fx**: fornix; **IFOF**: inferior fronto-occipital fasciculus; **ILF**: inferior longitudinal fasciculus; **PT**: pyramidal tract; **UF**: uncinate fasciculus. **(B-E)** Growth curves of DTI metrics. **(F-I)** Growth rates of DTI metrics. Lines represent the fitted mean curves (**B-E)** and their first derivatives (**F-I)**. For 95% simultaneous confidence bands, see Supplementary Fig. 5.

### Maturation of Infant White Matter Tracts Relative to Adults

To compare the diffusivity parameters of each infant tract to its mature adult stage, we used 40 adult diffusion data sets collected from the Human Connectome Project ^21^ and calculated the Mahalanobis distance between infant and adult tracts. Mahalanobis distance summarizes the distance between infant and adult tracts across multiple DTI measures (FA, AD, and RD) using a single metric. The Fx and the anterior thalamic radiation (ATR) were found to be among the most adult-like tracts, while the inferior longitudinal fasciculus (ILF), genu of the corpus callosum (CCg), arcuate fasciculus (AF), and inferior fronto-occipital fasciculus (IFOF) were among the least adult-like tracts (Fig. 3b-c). Changes in Mahalanobis distance between infant and adult tracts were nonlinear, with the largest (most negative) rate of change at birth, followed by rapidly slowing change rates until two months for most tracts (e.g., ILF, UF, IFOF, CCg) (Fig. 3d). A highly negative correlation was observed between the Mahalanobis distance of the white matter tracts and their change rates at birth (correlation=-0.9) to two months (correlation=-0.5) (Fig. 3e), indicating that less mature tracts may approach adult levels faster in early infancy. By 6 months, the Mahalanobis distance was less correlated with rate of change (correlation=-0.3), suggesting more uniformity among white matter tract development after 6 months.

**Figure 3.**
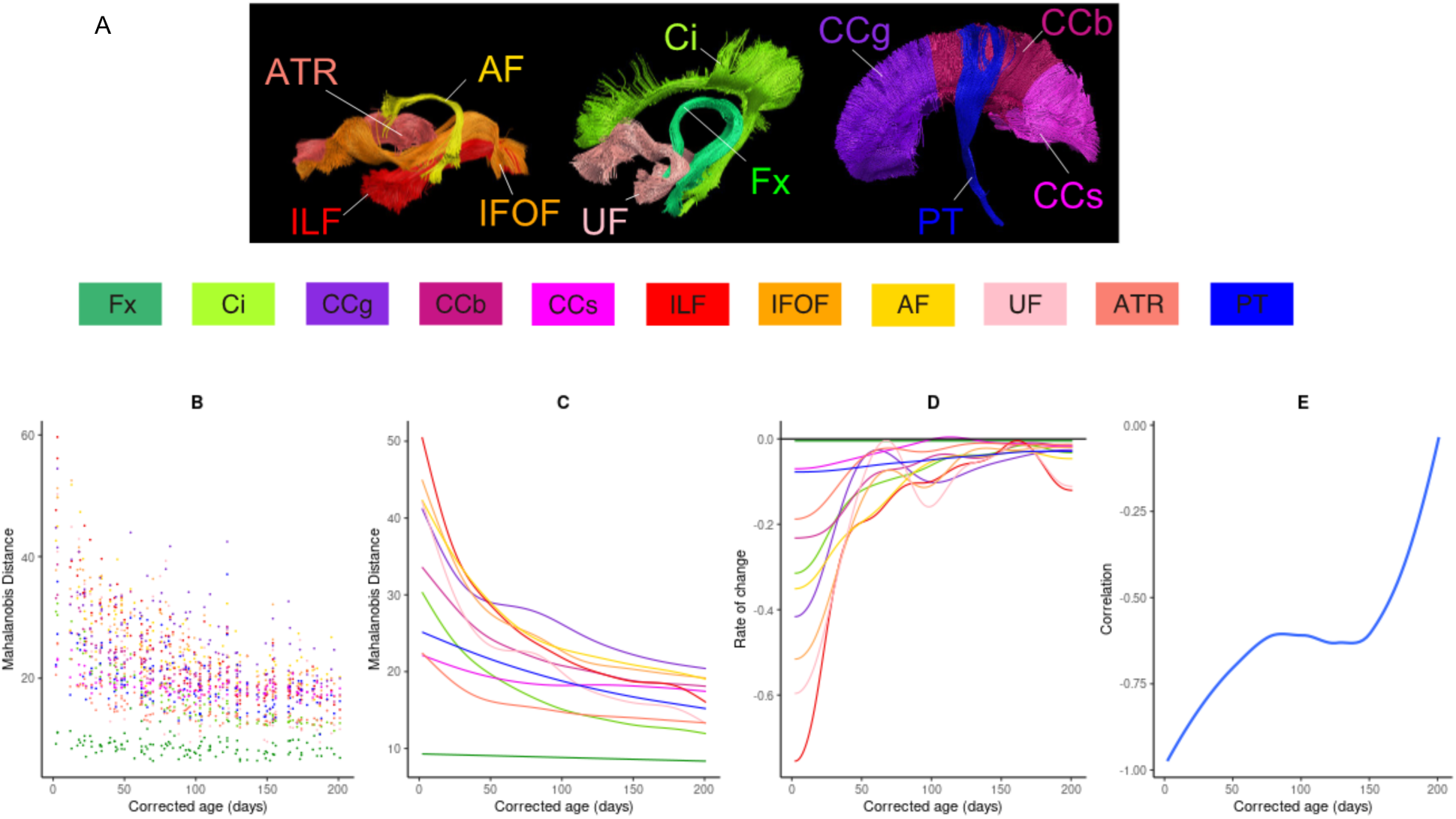
Mahalanobis distance between infant and adult white-matter pathways. (A) Delineation of 11 major white matter pathways. (**B**) Raw Mahalanobis distance measurements between infants and adults. (**C**) Fitted curves of Mahalanobis distance, based on FA, AD, and RD. Lower distances indicate more mature tracts. (**D**) Curves of rate of change of Mahalanobis distance. (**E**) Correlation between Mahalanobis distance and rate of change of the Mahalanobis distance (calculated from 11 fitted distances from each tract and the fitted rate of change at each corrected age). A highly negative correlation indicates distance is decreasing fastest in least developed tracts. For 95% simultaneous confidence bands, see Supplementary Fig. 6.

### Effects of Postnatal Experience on Rates of White Matter Maturation

We tested 3 hypotheses of the effect of gestational age at birth on white matter development. One possibility is that early exposure to the extrauterine environment may *accelerate* white matter maturation^26,27^. While accelerated maturation may be an adaptive response to the potential mismatch between a relatively immature white matter system and early exposure to the extrauterine environment, it may also enhance susceptibility to influences in early postnatal life. Accelerated maturation would be indicated by higher white matter change rates among infants with shorter compared to longer gestation, potentially resulting in a “catch up” effect among infants with shorter gestation when white matter maturation is expressed as a function of chronological age (time since birth), and even greater maturation among infants with shorter compared to longer gestation when white matter maturation is expressed as a function of corrected age (Fig. 4a).

**Figure 4.**
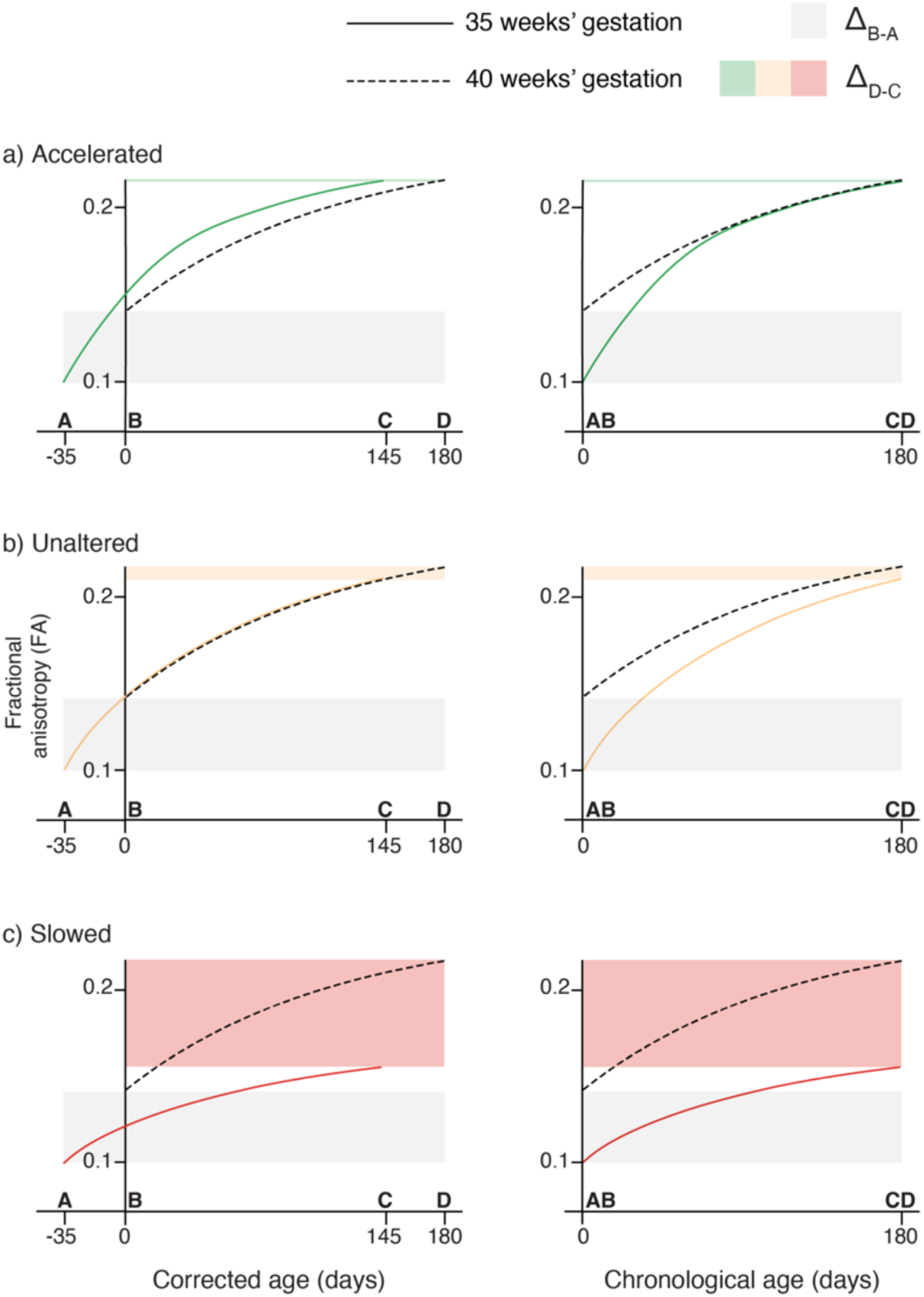
Hypothesized growth curves in an infant born at 35 weeks’ gestation (solid) and 40 weeks’ gestation (dashed) according to accelerated (a), unaltered (b) and slowed (c) hypotheses. A: corrected age at birth =-35 days, chronological age at birth = 0 days in infants born at 35 weeks’ gestation. B: corrected age at birth = 0 days, chronological age at birth = 0 days in infants born at 40 weeks’ gestation. C: corrected age = 145 days, chronological age = 180 days in infants born at 35 weeks’ gestation. D: corrected age = 180 days, chronological age = 180 days in infants born at 40 weeks’ gestation. The accelerated hypothesis predicts that infants born at 35 weeks’ gestation should ‘catch up’ to full-term (40 weeks’ gestation) infants when matched by chronological age and surpass fractional anisotropy (FA) maturation of full-term infants when matched by corrected age (FA at 145 days corrected age in 35 weeks’ gestation infant is equal to FA at 180 days in 40 weeks’ gestation infant, whereas FA is equal in 35 weeks’ and 40 weeks’ gestation infants at 180 days chronological age). The unaltered pace hypothesis predicts that FA in infants born at 35 weeks’ gestation and full-term infants should develop at the same rate when matched by corrected age. The slowed hypothesis predicts that FA in infants born at 35 weeks should develop at a slower rate, with differences in FA maturation becoming more pronounced over time.

A second possibility is that growth-rates of white matter development may be largely *unaltered* by early exposure to the extrauterine environment^28,29^. Unaltered maturation would predict similar white matter trajectories for infants with shorter and longer gestation when white matter maturation is expressed as a function of corrected age.

However, given reports of faster maturation in utero^30^, change rates would be faster when expressed as a function of chronological age, again potentially enhancing susceptibility (Fig. 4b).

Finally, a third possibility, based on recent evidence from ^30^, is that birth itself may *slow* trajectories of white matter development, potentially leading to a persistent reduction in white matter maturity amongst infants with shorter gestation (Fig. 4c). This slowed maturation would be indicated by reduced white matter maturation when expressed as a function of both corrected and chronological age.

### Effects of Gestational Age on White Matter Trajectories

We fit bivariate splines to examine the interaction between gestational age at birth and chronological age on FA, MD, AD, and RD. Our approach is summarized in Fig. 5 for the case of whole-brain FA. The 3D plot demonstrates that both chronological age and gestational age at birth increase FA (Fig. 5 a). The cross section of the bivariate spline at 35 weeks’ gestation and 40 weeks’ gestation is depicted in Fig. 5 b. We used simultaneous confidence bands to determine whether there was a significant difference between the cross section of the bivariate spline corresponding to 35 weeks’ gestation and 40 weeks’ gestation, where the red lines indicate the range over which the simultaneous confidence band of the difference curve does not contain zero (Fig. 5 c). We then plotted a dashed line corresponding to the maximum chronological age at which the curves significantly differed (Fig. 5 d).

**Figure 5.**
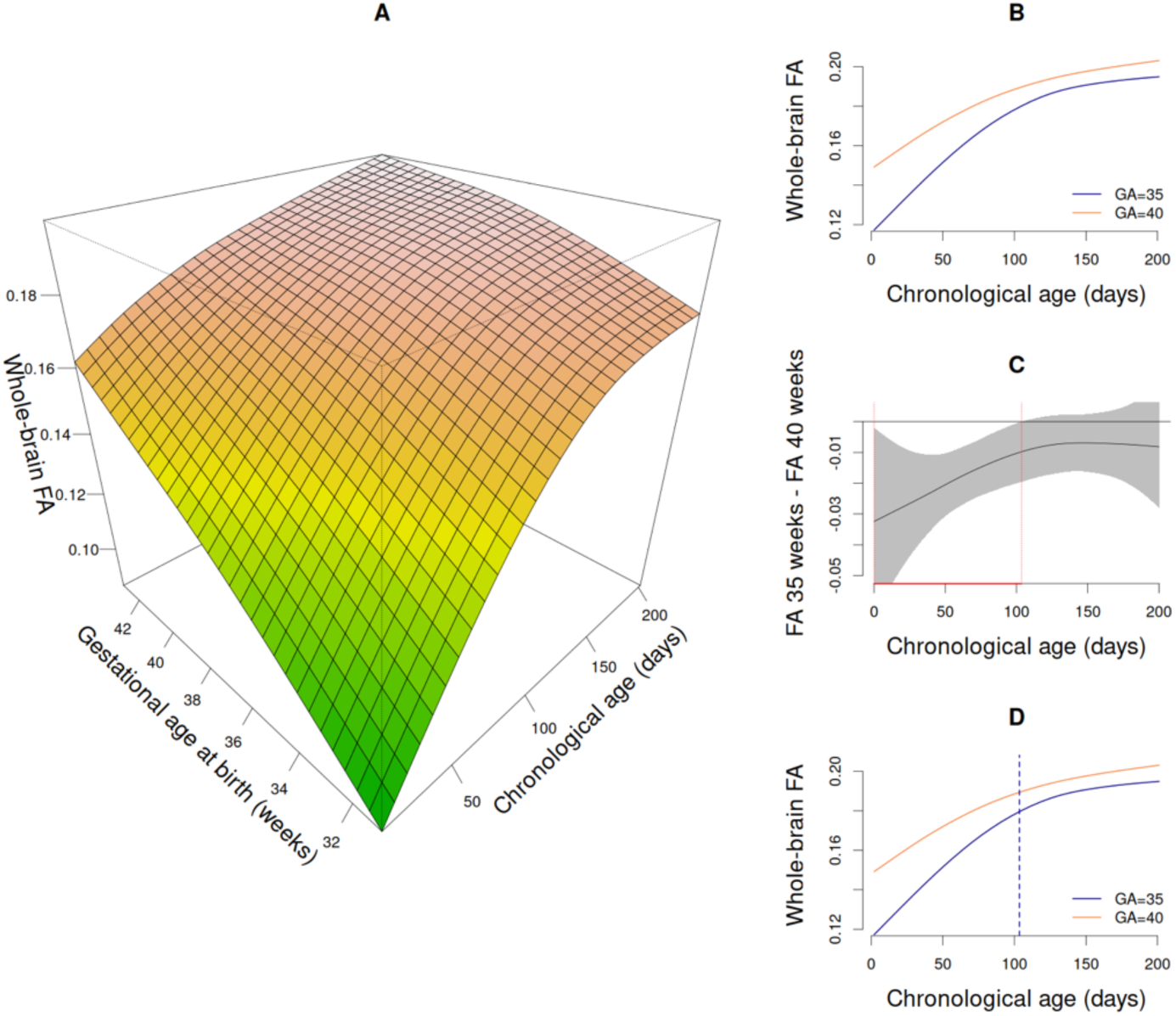
Illustration of the method used to characterize the impact of gestational age at birth (GA) and chronological age on infant brain white matter development. **(A)** 3D depiction of the bivariate spline estimating the effect of chronological age and GA on FA. (B) Growth curves at GA 35 weeks and GA 40 weeks extracted from the bivariate spline, which are generated by evaluating the bivariate spline at a fixed GA week while varying chronological age. **(C)** The difference between the GA 35 weeks and GA 40 weeks curves with 95% simultaneous confidence bands. Red lines indicate the range over which the difference curve is significantly different from zero. **(D)** The growth curves at GA 35 weeks and 40 weeks with the dashed vertical line corresponding to the maximum chronological age at which the simultaneous confidence band of the difference curve does not include zero.

Gestational age at birth had a significant effect on whole-brain white matter growth trajectories, with longer gestation associated with higher FA and lower MD, AD, and RD (i.e., more advanced maturation) (Fig. 6). In FA, the curve for infants born at 41 weeks is significantly higher than infants born at 40 weeks until 60 days and the curves for infants born at 39, 38, 37, 36, and 35 weeks are significantly lower than infants born at 40 weeks until 58, 69, 79, 91, and 103 days, respectively (Fig. 6 a). Similarly, the effect of gestational age at birth at 180 postnatal days tended to be weaker (flatter curve) than the effect of gestational age at birth at 30 postnatal days (Fig. 6 e), and growth curves were closer together as age increased (Fig. 6 e). This indicates an interaction between gestational age at birth and chronological age for FA, whereby the difference between infants born at younger gestational age and infants born at 40 weeks lessens over time. The effect of gestational age at birth at 180 postnatal days also tended to be weaker than the effect of gestational age at birth at 30 postnatal days in MD, AD, and RD (Fig. 6 f-h), but to a lesser extent than seen in FA (detailed difference curves are provided in Supplementary Fig. 7-10). In MD, the curves for infants born at 38, 37, 36, and 35 weeks are significantly lower than for infants born at 40 weeks until 60, 79, 98, and 157 days, respectively, while the curves for infants born at 39 and 41 weeks do not significantly differ from infants born at 40 weeks (Fig. 6 b). In AD, the curves for infants born at 37, 36, and 35 weeks have periods during which they significantly differ from infants born at 40 weeks, with no significant differences after 76, 95, and 154 days, respectively, while the curves for infants born at 38, 39, and 41 weeks do not significantly differ from infants born at 40 weeks (Fig. 6 c). In RD, the curves for infants born at 38, 37, 36, and 35 weeks are significantly lower than infants born at 40 weeks until 61, 79, 99, and 159 days, respectively, while the curves for infants born at 39 and 41 weeks do not significantly differ from infants born at 40 weeks (Fig. 6 d).

**Figure 6.**
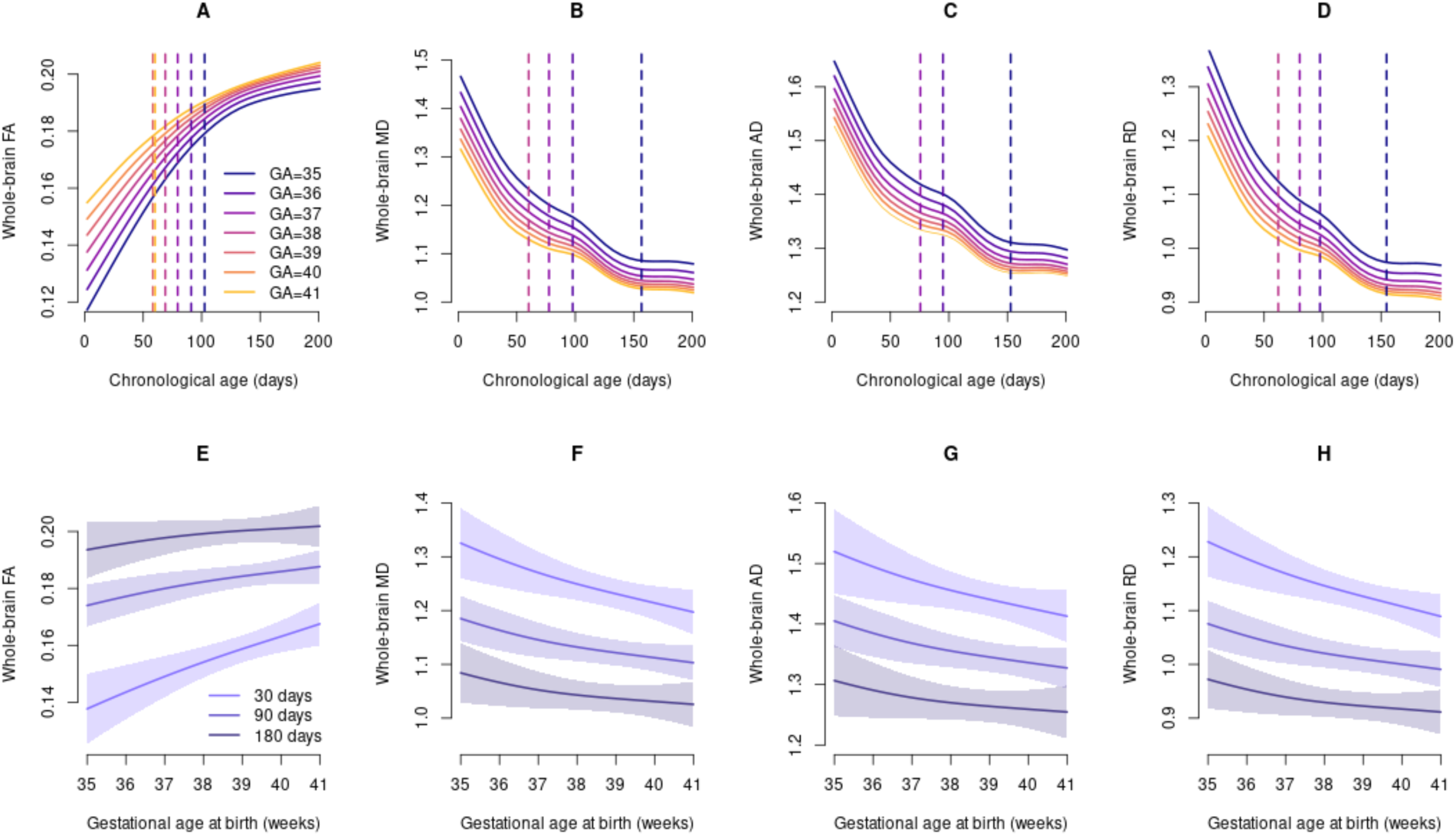
Effects of gestational age at birth (GA) on infant brain white matter development. (A)-(D) Growth curves of FA, MD, AD, and RD for infants’ born at different gestational ages (35 to 41 weeks). Dashed vertical lines denote the age (in postnatal days) when the simultaneous confidence bands for difference curves (between each GA curve and the curve for GA=40) contain zero. See Supplementary Fig. 7-10 for detailed plots of differences between GA. **(E)-(H)** The effects of GA at 30, 60, and 180 postnatal days for FA, MD, AD, and RD. Bands indicate 95% simultaneous confidence bands. The effect of GA is stronger (steeper curves) at 30 days; by 180 days, the effects of GA are minor.

These results, indicating that differences in white matter maturation between infants born at younger gestational ages and infants born at 40 weeks lessen over time, are inconsistent with the slowed hypothesis (see Fig. 4 c). To determine whether the observed effects of gestational age at birth on growth rates (i.e., initially reduced maturation and higher change rates among shorter gestation infants) reflect accelerated maturation amongst shorter gestation infants or unaltered maturation (see Fig. 4 a, b), we next constructed models using corrected age.

When corrected for gestational age at birth, the growth curves for children born at different gestational ages were very similar (Fig. 7). FA curves were nearly identical (Fig. 7a, Supplementary Fig. 11). There was a small trend towards a slowing of development in MD, AD, and RD after 100 days for infants born at earlier gestational age, but this change was not significant (Supplementary Fig. 12-14). This suggests that the higher change rates observed amongst infants with shorter gestation (Fig. 6) is likely consistent with the unaltered pace hypothesis, rather than accelerated maturation^26^.

**Figure 7.**
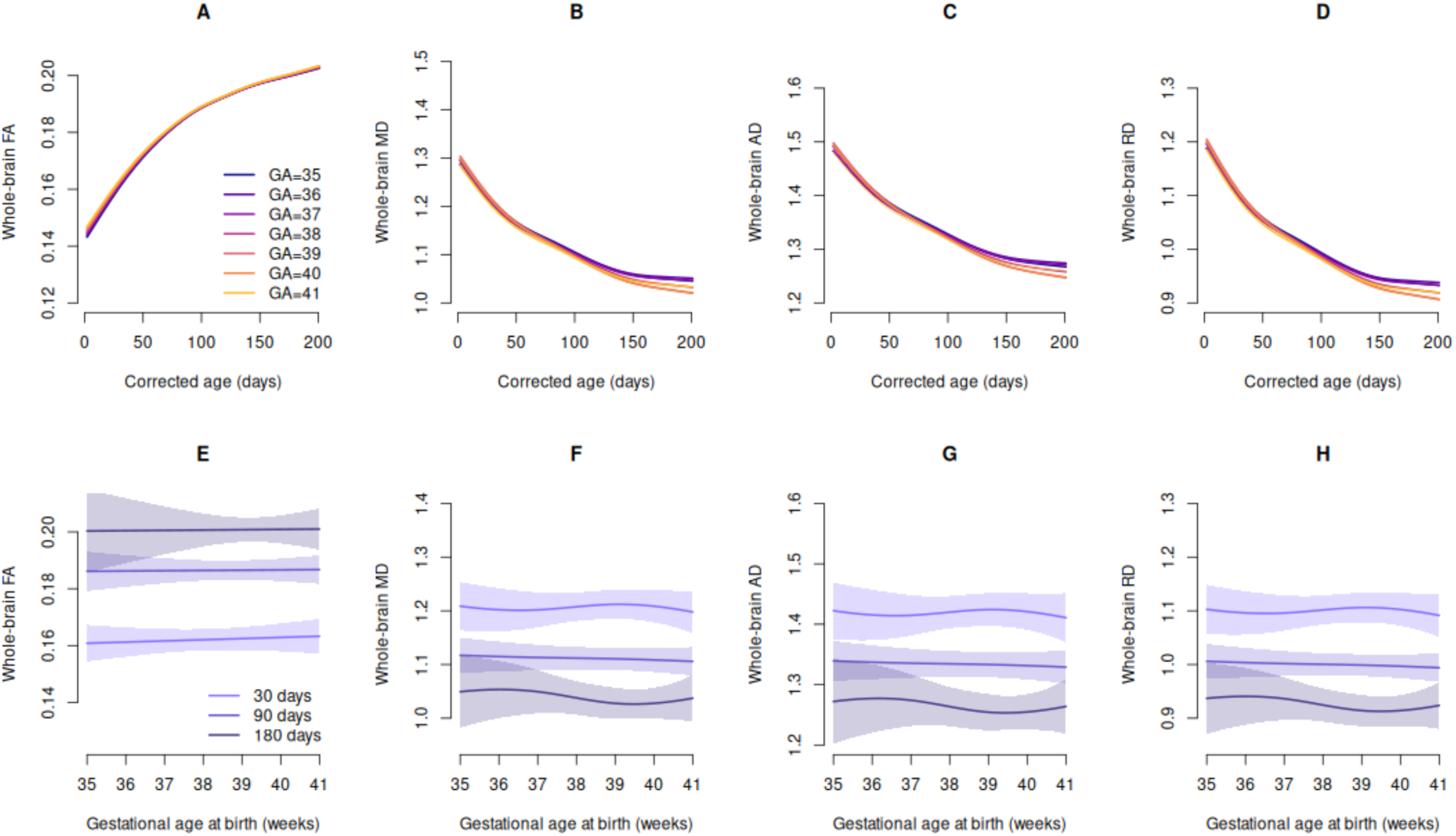
The effect of gestational age on growth curves when using corrected age. The effects of gestational age in models using corrected age are minor (see Supplementary Fig. 11-14 for difference curves in A-D plotted with 95% simultaneous confidence bands). In E through H, curves are nearly flat, indicating that gestational age at birth does not modify the developmental trajectories. The small slow down after 100 days in GA=35 does not result in significant differences from GA=40 (Supplementary Fig. 12-14).

We next examined whether the effects of gestational age at birth on FA were observed for all tracts. The FA growth curves at each gestational age became more similar with increasing chronological age for most tracts (Ci, CCg, CCb, CCs, IFOF, AF, UF, ATR, and PT, Supplementary Fig. 15). However, the rate at which tracts became more similar differed among tracts. For example, in the PT, FA for infants born at 35 weeks is significantly lower than infants born at 40 weeks until 65 days, while for CCb, FA for infants born at 35 weeks is significantly lower than infants born at 40 weeks until 126 days. ILF for infants born at 36 weeks and older was statistically indistinguishable from infants born at 40 weeks throughout the first six months (Supplementary Fig. 15). In MD, AD, and RD, in six tracts (Ci, CCb, CCs, ILF, IFOF, and UF), the differences between growth curves for different gestational ages decreased over time, but again occurring at different rates (Supplementary Figs. 16-18). For example, MD for infants born at 35 weeks differed from infants born at 40 weeks until 107 days in Ci, while MD differed until 143 days in IFOF. Finally, models constructed for each tract using corrected age showed no significant differences between age-corrected growth curves for infants born at different gestational ages (Supplementary Figs. 19-22).

### Effects of Sex on White Matter Trajectories

Whole-brain FA was initially higher in females than males, and did not significantly differ after 25 days, while whole-brain MD and RD were significantly lower until 10 and 14 days, respectively, while AD did not significantly differ (see Fig. 8 a-d).

**Figure 8.**
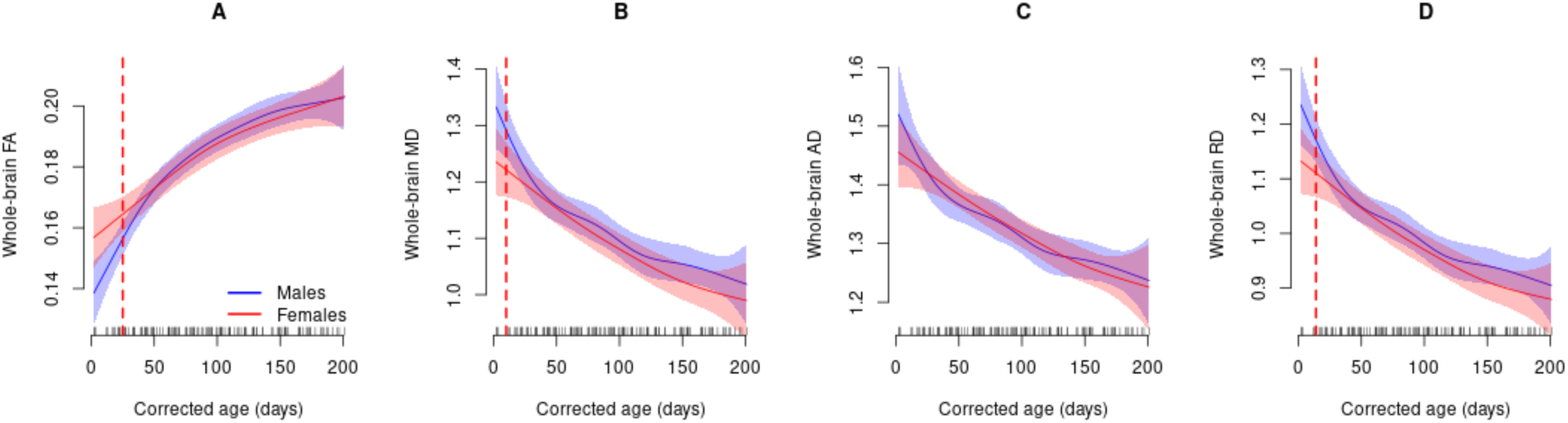
Whole-brain white matter growth curves for males (blue) and females (red). **(A)** Whole brain FA tended to be higher in females initially, and the simultaneous confidence band of the difference curve contained zero after 25 days (denoted by dashed line). **(B)** MD was initially higher in males, and the simultaneous confidence band contained zero after 10 days. **(C)** AD did not significantly differ between males and females. **(D)** RD was initially higher in males, and the simultaneous confidence band of the difference curve contained zero after 14 days.

## Discussion

Using one of the most densely sampled longitudinal pediatric neuroimaging datasets to date, we mapped temporally precise trajectories of white matter tract growth and growth rates from birth to 6 months of age. White matter development was highly dynamic and nonlinear, with growth rates of whole-brain white matter 6 to 11 times faster at birth than at 6 months of age. Regional growth trajectories and growth rates were asynchronous, with less mature tracts at birth developing faster postnatally. Gestational age at birth impacted white matter development, with shorter gestation infants having less mature tracts at birth, but faster growth rates, causing the white matter metrics to be more similar across different gestational ages at 6 months than at birth. Consistent with the unaltered pace hypothesis, white matter trajectories were similar when corrected for gestational age at birth.

### The First 3 Postnatal Months are a Dynamic Period of White Matter Development

While prior work has demonstrated that the most rapid postnatal changes in white matter development occur in infants’ first year and then slow thereafter^4,31,32^, our study is among the first to use finer temporal sampling (with scans covering the entire period between birth and 6 months) to more accurately model growth trajectories during early postnatal life. Change rates were highest at birth for FA, MD, AD, and RD (0.3%-0.4%/day) and substantially slower at 3 months (∼0.1%/day), continuing to slow to around 0.03%-0.05%/day after 5 months. These findings are consistent with a prior report that diffusion indices mature at a faster rate in the first 100 postnatal days compared to the rest of the year^33^, and may reflect underlying changes in aspects of white matter maturation, such as axonal organization and myelination, that are rapidly unfolding over this period. For instance, histological studies of parietal white matter in human infants show that phosphorylation of neurofilament markers (an index of axonal maturity) and expression of myelin basic protein (an index of myelination onset) increase rapidly between 37 and 54 postconceptional weeks (i.e., between birth and around 3 months)^34^. Striking changes in the concentrations of metabolites involved in axonal elongation and premyelination (NAA and glutamate, respectively) are also observed in the first 3 postnatal months compared to the rest of the first 2 years^35^.

Since the first 3 postnatal months are characterize by particularly rapid change in white matter tracts, exposure to deleterious or adaptive influences during this period could have large impacts on white matter trajectories and may be an important mechanism in the development of many conditions including Autism Spectrum Disorder, Schizophrenia and ADHD^3,4,35,36^. The temporally precise trajectories of white matter development described in the present study therefore provide crucial benchmarks for understanding how early departures from neurotypical trajectories—within specific pathways and at specific developmental time points—may yield distinct behavioral phenotypes.

### Regional Growth Trajectories and Dynamics of White Matter Development

White matter tracts had asynchronous maturational trajectories, suggesting that regional growth patterns may be governed by distinct biological mechanisms and may reflect adaptations to the infants’ pre-and postnatal environments ^37^. Analyses of the distance between infant and adult white matter tracts identified a limbic fiber, the Fx, and a projection fiber, the ATR, as being among the most mature, followed by the commissural fiber CCs and projection fiber PT. Association fibers, such as the ILF, AF, and IFOF, as well as the commissural fiber CCg, were among the least mature. These patterns are largely consistent with prior literature indicating that limbic and projection fibers tend to mature earlier than association fibers^38–42^.

Previous studies in older children and adults have shown that periods of significant change in white matter tract development coincide with behavioral changes ^43,44^. Hence, the longitudinal trajectories of major white development provided in the present study could provide an inroad for understanding how structural changes at the level of white matter are temporally related to the emergence and refinement of particular behaviors in early infancy. For instance, the first 2-3 postnatal months are a transformative period in infant homeostatic regulation and motor behaviors, with infants changing their feeding, sleeping, and waking patterns and acquiring new control of their eyes, neck, hands, and feet^45–53^. The relatively early maturation of limbic and projection fibers, such as the Fx and the PT (known to be involved in modulating physiological responses^54^ and motor behaviors^55^, respectively), may underlie (as well as be shaped by) these early changes in infant behavior.

We also observed differences in growth rates between tracts, with less mature tracts developing faster postnatally. This finding is consistent with the ‘speed-up’ hypothesis of white matter myelin development, advanced by ^56^, which proposes that white matter that is less myelinated at birth develops faster postnatally^56–58^. This hypothesis is contrasted with the ‘starts-first/finishes-first’ hypothesis, which posits that the pace of postnatal myelination follows earlier prenatal gradients of maturation^59^.

While the functional significance of the ‘speed-up hypothesis’ is not yet known, ^56^ proposed that the ‘catching up’ of less mature tracts to their more mature counterparts yields more uniform distribution of myelin across white matter, thereby allowing for more efficient and coordinated transmission of signals throughout the brain.

Asynchronous development across tracts (with some tracts maturing more rapidly than others), suggests that susceptibility may not be evenly distributed across tracts in the first 6 postnatal months. Less mature tracts at birth, such as the ILF, AF and IFOF, may be among the most susceptible during this period, since they are developing at higher growth rates. Consistent with this possibility, a study of periventricular leukomalacia patients found that developing tracts (e.g., the corona radiata and sagittal stratum) were more sensitive to injury than more developed tracts (e.g., limbic fibers)^60,61^. Given the disproportional involvement of association brain areas in functional specialization^62^ and neurodevelopmental-related conditions^63,64^, future research should investigate the relationship between the dynamics of white matter association tracts (which tend to be less mature and changing most rapidly) and their vulnerability in early infancy.

### Effect of Gestational Age at Birth on White Matter Growth Curves and Change Rates

This study is among the first to show that there is an association between gestational age at birth and growth curves and change rates of white matter development over the first postnatal months, even among near and full-term infants. When examined as a function of chronological age, we found that infants born at shorter gestation had lower FA and higher diffusivity values (i.e., less maturation), demonstrating that even minor differences in gestation are associated with differences in white matter. While this finding is perhaps unsurprising given the complex neurodevelopmental processes unfolding during late gestation, including neuronal migration, axon elongation, synaptogenesis, dendritic formation, and onset of myelination (see review by ^65^), it underscores the importance of considering the effect of gestational age at birth, even in studies of term-born infants.

Trajectories of white matter growth curves and change rates were consistent with the unaltered pace hypothesis. When matched on chronological age, infants with shorter gestation had higher FA change rates and, to a lesser extent, higher MD, AD, and RD change rates, potentially enhancing susceptibility in early postnatal life for infants born with shorter gestation. Correcting for gestational age at birth largely eliminated these differences in white matter growth curves and change rates between infants born at varying gestational ages.

The impact of gestational age on white matter was not uniform over infants’ first 6 months. When matched on chronological age, differences between infants with shorter gestation and infants born at 40 weeks’ gestation appeared to lessen over time. For infants born as little as 1 week early, it takes approximately 2 months for FA levels to be statistically indistinguishable from that of full-term (i.e., 40 weeks’ gestation) infants, with an additional 10 days required for each shortened week of gestation. The impact of gestational age at birth on white matter trajectories was also not uniform across tracts, with the effects of gestational age at birth on FA lessening earlier for some tracts (for example, the PT, where FA for infants born at 35 weeks is statistically indistinguishable from infants born full-term by 65 days) than others (for example, the CCb, where FA for infants born at 35 weeks is statistically indistinguishable from infants born full-term by 126 days). The trajectories presented herein therefore provide temporally precise, tract-specific windows of enhanced susceptibility for infants born at varying gestational ages, a critical step towards understanding the consequences of shortened gestational age on developmental outcomes.

Though our findings were consistent with the unaltered pace hypothesis, they do not preclude the possibility that gestational age at birth may alter the pace of other aspects of brain development or during particular time periods unexamined herein. For instance, Grotheer et al.^30^ found a slowing of white matter development very shortly after birth. Our findings do not eliminate the possibility that birth itself may result in an acute, transient period of slowing of white matter maturation in the first 1-2 postnatal weeks (where we only have four scans) that does not persist across the first 6 postnatal months. Additionally, given that our study focused on relatively large-scale, foundational aspects of structural brain architecture (i.e., the development of major white matter bundles), it is also possible that gestational age at birth may alter the pace of other aspects of brain circuity development. Indeed, several studies have shown that preterm infants have accelerated maturation of both stimulus-evoked^26,66^ and spontaneous functional brain activity across multiple sensory systems^27,67,68^, offering insight into the way in which functional brain systems may adapt to early exposure to the extrauterine environment. Finally, given that most of our sample was term-born, our findings may have captured subtler effects of gestational age at birth on white matter development compared to infants born very to moderately preterm (i.e., < 35 weeks’ gestation) who may show accelerated or slowed maturation as their relatively immature brain systems adapt to potential mismatches between their maturational state and environmental inputs and/or extended NICU stays that may more fundamentally alter early postnatal experience. Consistent with this possibility, a voxel-based morphometry study of preterm infants showed advanced white matter maturation in the sagittal striatum^69^.

### Sex effects

We found small effects of sex on white matter trajectories, with FA significantly higher in females than males during the first three weeks, and MD and RD significantly lower in females than males in the first week and two weeks, respectively, with no significant differences in AD. These findings are in line with previous reports of modestly advanced regionally specific white matter maturation in females compared to males from infancy to early childhood^42,58,70^. However, it is important to note that one study reported no sex differences in fiber pathways in neonates^71^, while others have observed more widespread sex differences in later life^72–76^, suggesting that white matter sexual dimorphism may be less apparent in early infancy and emerge in later life when exposure levels to sex hormones likely exert a greater influence on brain changes^70,72,73,77,78^.

### Limitations

While the DTI metrics used in this study (FA, MD, AD, RD) are sensitive to changes in white matter microstructure, such as axon diameter, spacing, orientation, and myelination, they do not allow for specific characterization of tissue microstructure. Other approaches, such as NODDI^79^, myelin water imaging^80^, or quantitative MRI measurements, such as longitudinal relaxation rate^56^, are required to disentangle the influence of specific microstructural properties of trajectories of white matter development. Additionally, FA and diffusivity measures can be less robust and reliable in curved structures with crossing fibers, like the AF and Fx^81,82^.

Finally, our sample of infant participants is predominantly White and from high socioeconomic status backgrounds and as such, is not representative of the racial and socioeconomic demographics of the city of Atlanta, Georgia, USA (where this study was based) (see Supplementary Table 1 for participant demographics).

### Conclusions

To summarize, this study provides the first longitudinal mapping of white matter development in infant’s first 6 postnatal months using densely sampled longitudinal data, providing temporally precise benchmarks of brain development, a necessary first step towards identifying, as early and precisely as possible, how departures from typical trajectories may lead to disability.

## Methods

### Design

Given that early infancy is marked by extremely rapid brain development^56,83^ and by individual differences in timing (when particular changes in growth occur) and in scale (how large or small a given change may be), dense longitudinal sampling during this period is critical to capture the regularities that characterize dynamic developmental processes^84,85^. Most published studies, however, have used cross-sectional^19,32,33,86–90^ or fixed longitudinal sampling^84,91,92^over the first years of life. While these studies contribute important knowledge about broad maturational changes from the first to the second year, dense longitudinal sampling, particularly in the first postnatal months—the period of most dramatic growth and change—is lacking. Additionally, most studies assume growth is linear ^83,87,88,90,93^ or make strong constraints on the shape of trajectories^84,91,94,95^, approaches that may miss crucial developmental spurts and arrests that would otherwise be captured by appropriate modeling^84,96^. Recent large-scale studies have used more advanced approaches to model brain morphometry and white matter changes across the lifespan, but studies modeling early life changes in white matter maturation, particularly during 0 to 6 months, are lacking ^97–99^. Consequently, growth and growth-rate trajectories of white matter development during the first months after birth—a dynamic period of growth that may represent a sensitive period for susceptibility in neurodevelopmental disorders—remain largely uncharted.

As noted in the main text, in the present study, diffusion MRI data were collected using a non-uniform longitudinal sampling design, with data collected from each infant at up to 3 randomized time points between birth and 6 months from 79 typically developing infants (see Supplementary Table 1 for participant demographics). This approach yielded dense coverage over the 0-to 6-month period (129 scans, each separated by a mean of only 1.55 days (stdev=1.69, min=0 days, max=9 days), a necessity for generating temporally precise trajectories during periods of rapid developmental change^20^. See Supplementary Fig.1 for a distribution of participant age at each scan.

### Participants

Participants were typically developing infants (48 male, 31 female) enrolled in an ongoing prospective longitudinal study of infants at low-and elevated-likelihood for Autism Spectrum Disorder (ASD). Typicality was ascertained by the absence of factors associated with increased likelihood of developmental disability: all infants had uncomplicated deliveries (mean gestational age = 38.6 weeks, SD=1.98 weeks, 87.3% of infants born at ≥ 37 weeks’ gestation), had no family history of ASD in first, second, or third degree relatives, no developmental delays in first degree relatives, no pre-or perinatal complications, no history of seizures, no known medical conditions or genetic disorders, and no hearing loss or visual impairment. Participants had no contraindications for MRI. The research protocol was approved by the Emory University Institutional Review Board.

All infants were invited to complete 3 scans between birth and 6 months. Diffusion MRI data were collected from 81 infants with a total of 135 scans ranging between 10 and 210 postnatal days. Infants awoke before a full diffusion data set was collected in 16 out of 135 scans. Incomplete scans were excluded if less than 6 volumes of diffusion weighted images (the minimum number of volumes required for estimating diffusion tensor and tensor-based metrics) were collected^100^ (n=4) or if eddy current distortion correction could not be performed (n=2). The final data set consisted of 129 scans collected from 79 infants, with 55.7%, 32.9% and 11.4% of infants contributing one, two and three longitudinal scans, respectively.

### MRI data acquisition

All infant scans were acquired at Emory University’s Center for Systems Imaging Core on a 3T Siemens Tim Trio (n=52 scans) or a 3T Siemens PrismaFit (n=77 scans) scanner, using a 32-channel head coil. All infants were scanned during natural sleep, using procedures similar to those described in^101^ (see Supplementary Materials for details).

Diffusion MRI data from the Tim Trio scanner were acquired using a multiband sequence with the following parameters: repetition time (TR) of 6200 ms, echo time (TE) of 74 ms, a multiband factor of 2 combined with parallel imaging (GRAPPA) with an acceleration factor of 2, a field-of-view (FOV) of 184×184, image matrix of 92×92, b value of 700 s/mm^2^, spatial resolution of 2 mm isotropic, and 61 diffusion directions, 56 slices covering the whole brain. Six b=0 volumes were collected to improve the signal-to-noise ratio (SNR) of the baseline diffusion MRI signal^102^. The total scan time for the diffusion MRI sequence was 7 minutes 26 seconds.

Diffusion MRI data from the Prisma Fit scanner were acquired using a multiband sequence with the following parameters: repetition time (TR) of 2330 ms, echo time (TE) of 86.6 ms, a multiband factor of 4 without parallel imaging acceleration, a field-of-view (FOV) of 184×184, image matrix of 106×106, b value of 700 s/mm^2^ with 6 b=0 volumes, spatial resolution of 1.75mm isotropic, and 89 diffusion directions, 68 slices covering the whole brain. The total scan time for the diffusion MRI sequence was 3 minutes 58 seconds.

*Adult Data.* To compare the diffusivity parameters of each infant tract to its mature adult stage, we used 40 adult diffusion data sets collected from the HCP (https://www.humanconnectome.org) ^21,102,103^. HCP data were collected on a Skyra 3T scanner using a multiband sequence with the following parameters: repetition time (TR) of 5520 ms, echo time (TE) of 89.5 ms, a multiband factor of 3, a field-of-view (FOV) of 210×180, image matrix of 168×144, b value of 1000 s/mm^2^, spatial resolution of 1.25 mm isotropic and 90 diffusion directions. The first shell was utilized as the reference.

Additional data acquisition details can be found in^21,104^.

### Data preprocessing

Infant data were preprocessed using FSL (6.0.03) and in-house MATLAB (R2016b, MathWorks Inc.) code. Preprocessing steps included correcting inter-volume motion artifacts, removing susceptibility distortion using the “topup” function in FSL, and eddy-current distortion and motion correction using FSL’s “eddy” tool^105^. Diffusion MRI parameters were estimated using weighted least squares ^106,107^.

Adult data were preprocessed using standard HCP pipelines (please see^21^ for details).

### Image Registration

Tensor-based registration was used to align infant brain images to a common space. Unlike T1-and T2-weighted images which are isointense by approximately 6 months of age^108^, tensor maps maintain gray and white matter signal intensity differences over the first postnatal months and also provide orientation information about white matter microstructure, thereby allowing for more detailed mapping of features between individuals^109^. Infant brain images were aligned to a cohort-specific template using multilevel registration^110,111^. See Supplementary Information for more details. The registered tensor maps were then averaged, creating an average tensor map with high SNR. The same approach was used to align adult images to a common space.

#### Fiber Tractography

##### Infant white matter pathways

Whole-brain tractography was seeded from the average whole-brain white matter voxels (mean fractional anisotropy, FA >0.15) using FACT^112,113^ in Camino ( http://camino.cs.ucl.ac.uk/). After all possible streamlines representing white matter connectivity in the infant brain were constructed, 11 major white matter pathways were delineated using TrackVis (http://www.trackvis.org), using methods described in previous literature (see Supplementary Fig. 2 for details). Tracts of interest included the arcuate fasciculus (AF), anterior thalamic radiation (ATR), cingulum (Ci), fornix (Fx), inferior fronto-occipital fasciculus (IFOF), inferior longitudinal fasciculus (ILF), uncinate fasciculus (UF), pyramidal tract (PT), genu of the corpus callosum (CCg), body of the corpus callosum (CCb), and splenium of the corpus callosum (CCs). The whole-brain mask was created by combining all tracts and white matter of the corpus callosum and binarizing them to form a single mask. Two-dimensional views of the tracts are shown in Figs. 2 and 3 and three-dimensional views are shown in Supplementary Fig. 3.

After delineating the tracts in the cohort-specific template space, tracts were propagated to each scan space using the transformation maps from the tensor-based registration. This two-step approach allowed us to control variability due to the differential traceability of tracts across different ages.

##### Adult white matter pathways

To compare infant white matter tracts to adults, the infant cohort-specific template was registered to the HCP adult template using tensor-based registration, and then this map was used to register the infant white matter tract masks to the adult space. This ensures that any differences in DTI metrics observed between infants and adults are likely due to microscopic, rather than macroscopic, differences.

##### DTI-derived metrics for quantifying white matter development

Fractional anisotropy (FA), mean diffusivity (MD), axial diffusivity (AD), and radial diffusivity (RD) were used to quantify tract-specific white matter development^39,100,114^. For each scan, mean FA, MD, AD, and RD values were calculated within each tract-of-interest and averaged across hemispheres (for all tracts, with the exception of the corpus callosum).

##### Harmonization of data collected on Siemens Trio and PrismaFit scanners using Longitudinal ComBat

We used Longitudinal ComBat to harmonize infant data collected on Siemens Trio and PrismaFit 3T scanners. Longitudinal combat is a harmonization technique that models additive and multiplicative site/scanner effects while accounting for the correlation between repeated measurements on the same participant^115^. Data harmonization was performed separately for each DTI metric and each tract using the “longCombat” R package. We included scanner as the batch effect, participant ID as a random effect, and sex and corrected age as main effects. For corrected age, we accounted for possible non-linearities using a cubic b-spline with knots at the 33.3 and 66.7 percentiles. After harmonization, the effect of scanner was not significant (see Supplementary Figs. 23-26 for details), suggesting the effectiveness of Longitudinal ComBat in reducing unwanted scanner effects. Therefore, the covariate scanner was not included in subsequent regression analyses.

##### Growth Trajectory Analysis Using Generalized Additive Mixed Effect Models (GAMMs)

GAMMs were used to estimate growth trajectories from longitudinal measures of infant DTI-derived metrics^116–118^. This approach does not assume any fixed form between predictors and responses (e.g. linear or quadratic), but instead identifies best-fit trajectories in a data-driven fashion. Whole-brain growth curves (Fig. 1 A-D) and tract-specific growth curves (Fig. 2) were estimated using a smooth for corrected age, defined as the age in days minus 7*(40-gestational age in weeks), and a random intercept for infant. These models used the default spline smoother (thin-plate regression splines) with the default degrees of freedom, and we used restricted maximum likelihood to determine the amount of smoothing, implemented in the “mgcv” R package^118^. We assessed the adequacy of the spline DFs by comparing the effective degrees of freedom to the maximum possible (9), where the default was considered sufficient when the edf was not close to 9, here defined as less than 6. Rates of growth (Fig. 1 e-h) were estimated by calculating the first derivative of the GAMM-fitted growth trajectory using the “gratia” R package.

We calculated the Mahalanobis distance for each infant *i*, scan *j*, and tract *k* as the distance between the vector of infant DTI measures, denoted **x**_ijk_ =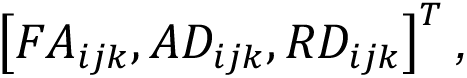 and the mean of adult space, denoted 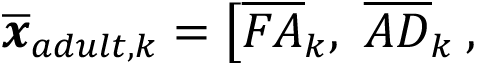 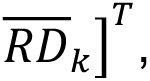 normalized by the covariance of FA, AD, and RD in the adult sample, denoted Σ_adult,k_ The Mahalanobis distance is

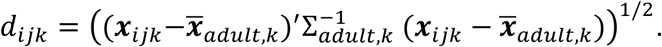

MD was not included in the vector of DTI measures because it is a linear combination of AD and RD, such that the Mahalonobis distance would be undefined. Mahalanobis distance curves (Fig. 3) were estimated using a smooth for corrected age and a random intercept for infant, following the same approach as the whole-brain and tract-specific GAMMs described above. Using the predicted distances for each of the *k* tracts at corrected age *t*, denoted 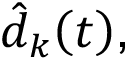 and the estimated rate of change of the Mahalanobis distances, denoted 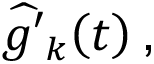 we calculated the correlation between the Mahalanobis distance and the rate of change, i.e., 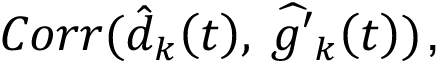 where the correlation was calculated using the 11 data points corresponding to the tracts. This was repeated for *t*=2 to 201, and the correlations were then visualized using a loess smoother with default settings implemented in the R package “ggplot2”^119^.

To examine the interaction between gestational age and chronological age on DTI metrics (Fig. 5, 6), we fit GAMMs using a tensor spline for gestational age and chronological age with a main effect for sex and a random intercept for infant as described in the Supplementary Information. Predicted curves and simultaneous confidence bands were calculated using the “itsadug”^120^ R package with sex set to male. To examine the interaction between sex and corrected age (Fig. 8 a-c), we specified sex as the “by” variable in the spline smooth for corrected age. Details of model fitting are provided in Supplementary Information.

## Acknowledgments

We greatly appreciate the families and their infants who volunteered to participate in this research study. We would also like to thank the research coordinators, assistants, and fellows at the Marcus Autism Center, Brittney Sholar, Carly Reineri, Joannna Beugnon, Lindsey Evans, Jordan Pincus, Jennifer Gutierrez, Tristan Ponzo, and Adriana Mendez, and the MRI Techs at the Emory Center for Systems Imaging Core, Michael White, Sarah Basadre, and Samira Yeboah, for their data collection efforts, as well as Dr. Lei Zhou and Michael Valente for their assistance with equipment development and data acquisition protocols, and Mahmoud Zeydabadinezhad for his help with data preprocessing. Finally, we would like to thank Dr. Sasha Greenspan for her assistance with Figure 4, and Dr. Aiden Ford for discussion and suggestions.

## Author Contributions

B.R.: formal analysis, software, validation, visualization, writing-original draft; L.L.: methodology, formal analysis, validation, visualization, funding acquisition, writing-review & editing; W.J: conceptualization, funding acquisition, writing-review & editing; S.S.: conceptualization, investigation, data curation, visualization, supervision, project administration, resources, funding acquisition, writing-original draft.

## Data Availability

Data collected from National Institute of Mental Health (NIMH) 2P50MH100029, R01MH118285, and R01MH119251 are available from the NIMH Data Archive (NDA). NDA is a collaborative informatics system created by the National Institutes of Health to provide a national resource to support and accelerate research in mental health. Code is available at https://github.com/thebrisklab/WhiteMatterDevelopment. Data will be released in a NDA study-specific repository upon publication. The manuscript reflects the views of the authors and may not reflect the opinions or views of the NIH.

