## Supplemental Materials for "Dynamics of infant white matter maturation from birth to 6 months"

*MRI Data Acquisition Procedure*

Infant scans were acquired at Emory University's Center for Systems Imaging Core on a 3T Siemens Tim Trio (n=52) or a 3T Siemens Prisma (n=77) scanner, using a 32-channel head coil. All infants were scanned during natural sleep, using the following procedure. First, infants were swaddled, rocked, and/or fed to encourage natural sleep. Once asleep, the infant was placed on a pediatric scanner bed. Scanner noise was reduced below 80 dBA by using: 1) sound attenuating pediatric headphones, equipped with MR-safe optical microphones to enable real-time monitoring of in-ear sound levels throughout the scan session; and 2) a custom-built acoustic hood, inserted into the MRI bore. To mask the onset of scanner noise, white noise—gradually increasing in volume—was played through the headphones prior to the first sequence. An MRI-compatible camera (MRC Systems) was mounted on the head coil to enable monitoring of the infant throughout the scan. A trained experimenter remained in the scanner room and the procedure was stopped if the infant awoke or if an increase in sound level was observed.

*Image Registration*

Infant brain images were aligned to a sample-specific template using multilevel registration [1,2] First, each infant's tensor map was aligned to the tensor map of a randomly chosen infant participant using 6-degree of freedom (df) rigid body transformations. Second, the aligned images from all participants were averaged to create the initial 6-df target template. Third, individual tensor maps were aligned to the initial 6-df target template using 12-df affine transformations and then averaged to form the intermediate 12-df target template. Fourth, individual tensor maps were registered to the 12-df intermediate target template using diffeomorphic registration and then averaged to create the sample-specific diffusion tensor template[1]. After the sample-specific diffeomorphic template was built, individual tensor maps were aligned to the template space using rigid (i.e., 6-df), then affine (i.e., 12-df), and then diffeomorphic registration, as described above for group analysis.

*Effects of Scanner*

After harmonizing the data collected on Siemens Trio and Prisma scanners using longitudinal Combat[3], we examined the effect of scanner (Trio vs Prisma) on the harmonized DTI metrics:

$$DTI_{ij} = \beta_0 + \beta_1 male_i + \beta_2 Trio_i + f_1\{AgeScanC_{ij} * (1 - Trio_i)\} + f_2(AgeScanC_{ij} * Trio_i) + b_i + \varepsilon_{ij},$$

where $DTI_{ij}$ denotes the whole-brain DTI metric (FA, MD, RD, or AD) for the *i*th infant at time point *j*; $male_i$ is an indicator variable equal to one if the child is male; $Trio_i$ is an indicator variable equal to one if the child was scanned using the Trio; $AgeScanC_{ij}$ is the gestationally corrected age in days for the *i*th infant at the *j*th MRI scan; $b_i \sim N(0, \tau_2)$ is a normally distributed random intercept for child; and $\epsilon_{ij} \sim N(0, \sigma^2)$ is the error. Next, $\beta_0$, $\beta_1$, and $\beta_2$ are parametric terms for the intercept, male, and Trio, and $f_1\{AgeScanC_{ij} * (1 - Trio_i)\}$ is a smooth function of gestationally corrected age in days for children scanned on the PrismaFit, and $f_1\{AgeScanC_{ij} * (1 - Trio_i)\}$ is a smooth function for children scanned on the Trio. The effect of scanner was not significant (Supplementary Figs. 15-18), suggesting the effectiveness of longitudinal Combat in reducing unwanted scanner effects. Therefore, the covariate scanner was not included in subsequent regression analyses.

*Gestational Age Effect (Fig. 6 and 7)*

The effect of gestational age at birth was examined in the following model:

$$DTI_{ij} = \beta_0 + \beta_1 male_i + f_1(AgeScan_{ij}) + f_2(GestationalAge_i) + f_{12}(AgeScan_{ij}, GestationalAge_i) + b_i + \varepsilon_{ij},$$

where $AgeScan_{ij}$ is the (uncorrected) age at scan in days, $GestationalAge_i$ is the gestational age at birth for the *i*th infant, and other terms were previously defined. Here, $f_{12}(AgeScan_{ij}, GestationalAge_i)$ is a tensor spline constructed in the null space of the splines from $f_1(AgeScan_{ij})$ and $f_2(GestationalAge_i)$ using the function "ti()" in the "mgcv" R package. This bivariate spline allows different growth trajectories for different gestational ages. The tensor spline default is k=5 for each dimension. In FA, this resulted in an effective degrees of freedom (edf) close to the maximum for $f_1(AgeScan_{ij})$, and hence we re-fit the model with "k=9" for $f_1$, "k=5" for $f_2$, and "k=c(9,5)" for $f_{12}$, which resulted in edf's well below their maxima (edf=3.5, 2.0, and 2.1 for $f_1$, $f_2$, and $f_{12}$, versus maximums of 8, 4, and 32). These specifications were also used for MD, AD, and RD.

After correcting the chronological age using gestational age (i.e., corrected age, which is defined as chronological age +/- the number of weeks over/under 40 weeks), the differences at different gestational ages are minor (Supplementary Fig. 19). Corrected age was included while gestational age was not included in sex-specific subsequent models.

*Sex Effects (Fig. 8)*

Sex effects were examined using the following model:

$$DTI_{ij} = \beta_0 + \beta_1 male_i + f_1\{CorrAgeScan_{ij} * (1 - male_i)\} + f_2(CorrAgeScan_{ij} * male_i) + b_i + \varepsilon_{ij},$$

$f_1\{CorrAgeScan_{ij} * (1 - male_i)\}$ is a smooth function of corrected age for the *i*th infant at the *j*th MRI scan if the infant is female, and $f_2(CorrAgeScan_{ij} * male_i)$ is the smooth function for males.

| | (N = 79) |
|---|---|
| **Infant Sex** | 31f, 48m |
| **Gestational Age at birth** | |
| *Very Preterm (< 32 weeks)* | 2.5% |
| *Late Preterm (34-36 weeks)* | 10.1% |
| *Early Term (37-38 weeks)* | 22.8% |
| *Full Term (39-40 weeks)* | 55.7% |
| *Late Term (> 41 weeks)* | 8.9% |
| **Race (N=68)** | |
| *Black* | 8.8% |
| *Native* | 1.5% |
| *White* | 86.8% |
| *More than one race* | 2.9% |
| **Maternal Education (N=67)** | |
| *High School* | 1.5% |
| *College Courses* | 6.0% |
| *Associate's Degree* | 1.5% |
| *College Degree* | 32.8% |
| *Graduate Degree* | 58.2% |
| **Household Income (N=65)** | |
| *< $40000* | 7.7% |
| *$40000-$80000* | 16.9% |
| *$80001-$100000* | 18.5% |
| *$100001-$150000* | 26.2% |
| *> $150000* | 30.8% |

**Table S1: Demographics of participant sample.** Nine participants did not complete the Family Demographic Form, and additional participants declined to answer specific questions about Race, Maternal Education, and Household Income. For information categories with a participant number less than the total sample, the N is specified next to the category title.

### Figures

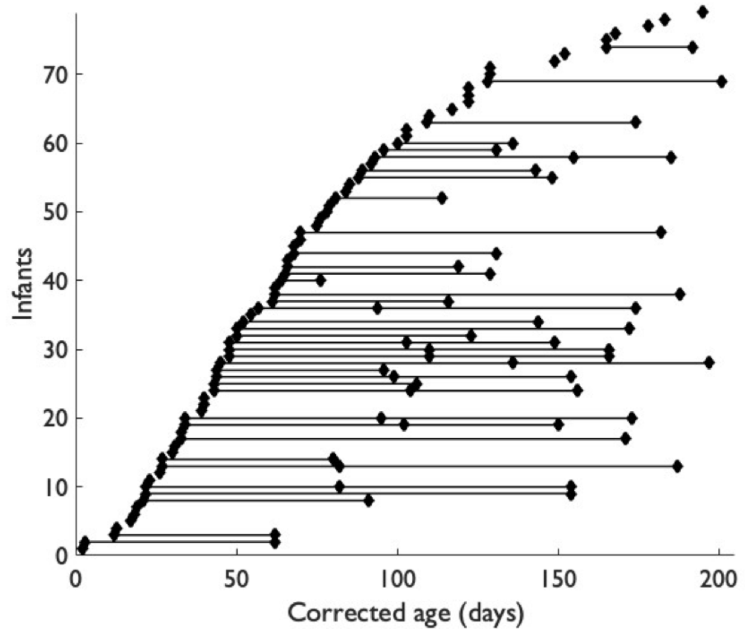


**Supplementary Figure 1**. **The distribution of scans by corrected chronological ages for all included participants.** Each dot represents one diffusion MRI scan from a participant and dots connected by lines represent all available longitudinal scans from that given participant. Data were collected using a non-uniform longitudinal sampling design, with diffusion MRI scans collected from each infant at up to 3 randomized time points between birth and 6 months (129 scans total, each separated by a mean of 1.55 days (stdev=1.69, min=0 days, max=9 days).

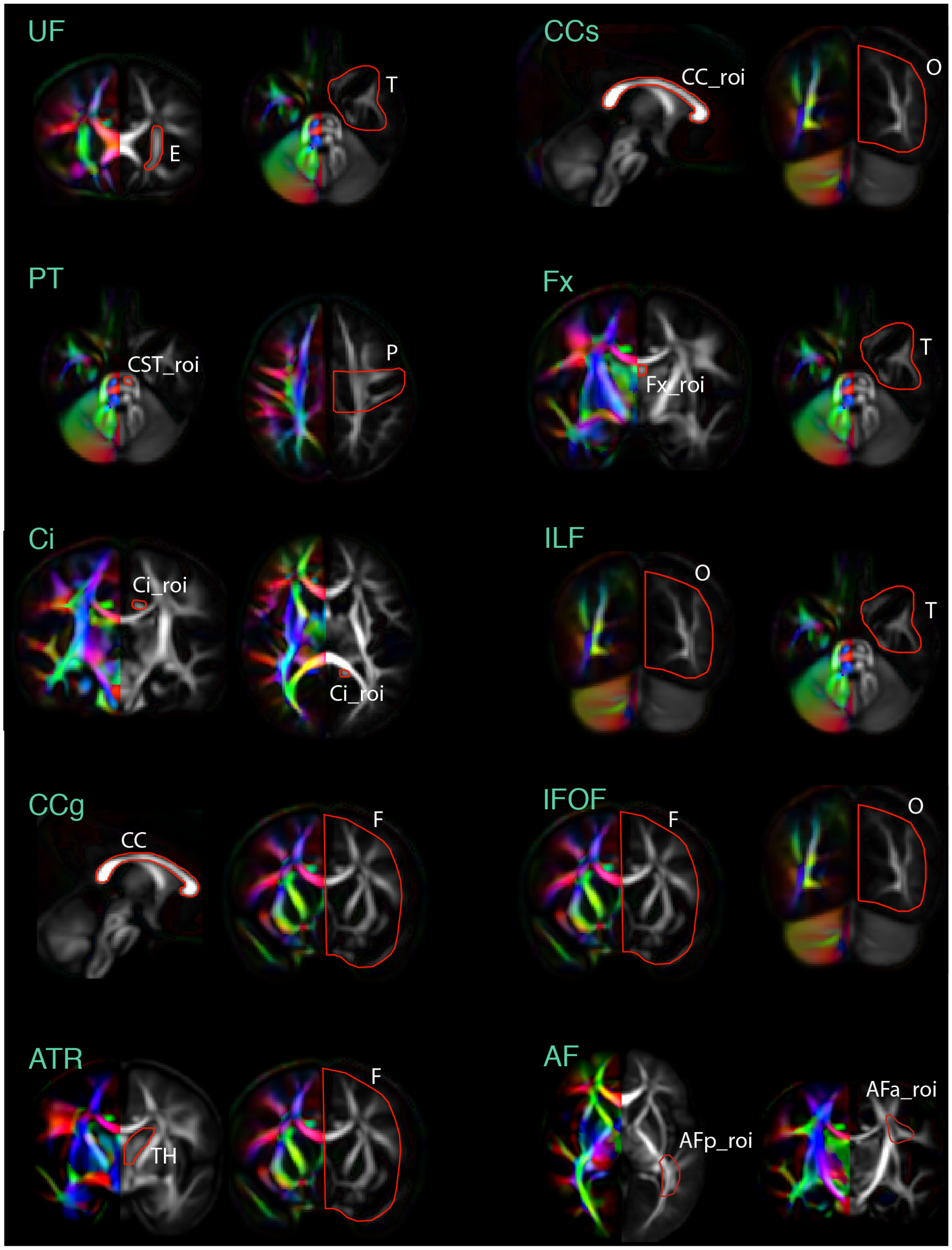


**Supplementary Figure 2. Defining regions of interest (ROIs) for delineating 11 major white matter pathways in infant brains.** The method for delineating these tracts largely follow those outlined by Catani et al.[4] and others[5-9]. Names of delineated white-matter tracts are annotated in green. The ROIs used to delineate these tracts are annotated in white. For delineated tracts, **AF**: arcuate fasciculus; **ATR**: anterior thalamic radiation; **CCg**: genu of corpus callosum; **CCs**: splenium of corpus callosum; **Ci**: cingulum; **Fx**: fornix; **IFOF**: inferior fronto-occipital fasciculus; **ILF**: inferior longitudinal fasciculus; **PT**: pyramidal tract; **UF**: uncinate fasciculus. For the ROIs used to delineate tracts, **E**: external capsule ROI; **T**: temporal ROI; **CC_roi**: corpus callosum ROI; **O**:

occipital ROI; **CST_roi**: corticospinal tract ROI; **P**: pre- and post-central gyrus ROI; **Fx_roi**: fornix ROI; **Ci_roi**: cingulum ROI; **F**: frontal ROI; **TH**: thalamus ROI; **AFp_roi**: posterior arcuate fasciculus ROI; **AFa_roi**: anterior arcuate fasciculus ROI. UF, AF, ILF, and IFOF are association fibers; Ci and Fx are limbic fibers; CCg, CCb, CCs, are commissural fibers; and PT and ATR are projection fibers.

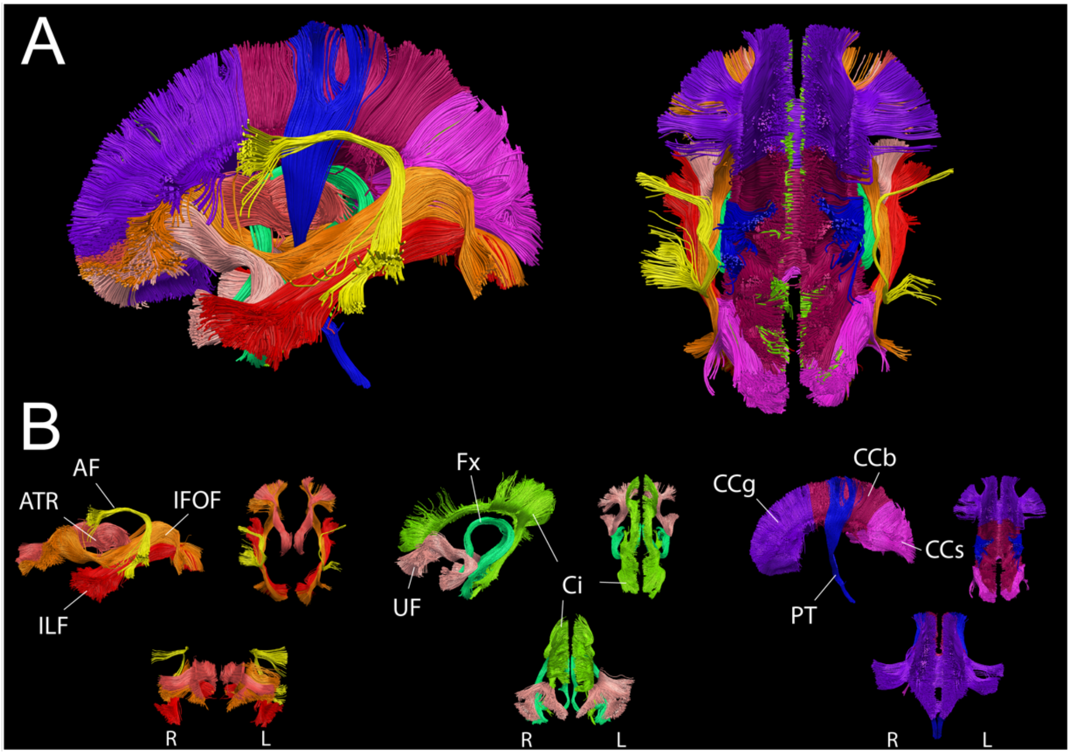


**Supplementary Figure 3: Delineation of 11 major white matter tracts. AF**: arcuate fasciculus; **ATR**: anterior thalamic radiation; **CCb**: body of corpus callosum**; CCg**: genu of corpus callosum; **CCs**: splenium of corpus callosum**; Ci**: cingulum; **Fx**: fornix; **IFOF**: inferior fronto-occipital fasciculus; **ILF**: inferior longitudinal fasciculus; **PT:** pyramidal tract; and **UF**: uncinate fasciculus.

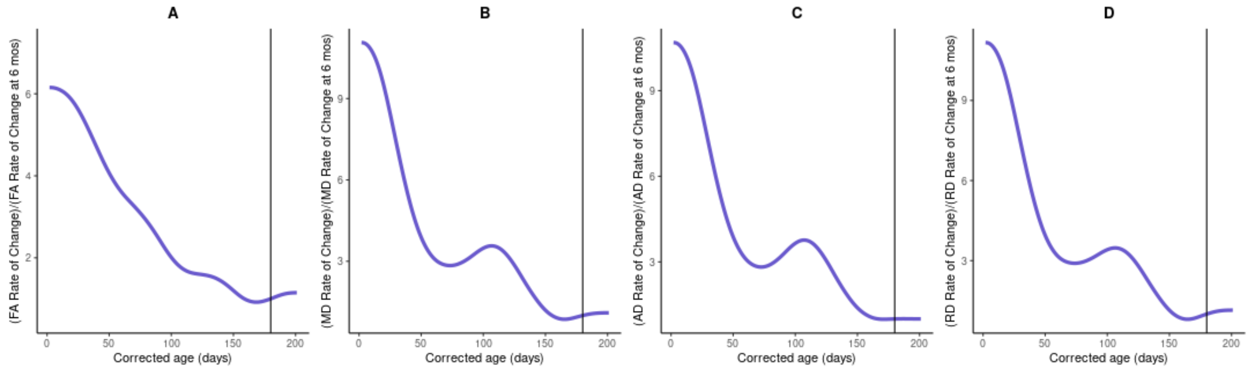


**Supplementary Figure 4: Number of folds of growth rates in whole-brain white matter relative to the growth rate at 6 months for all 4 DTI metrics.** Growth rates normalized to the growth rate on day 180 (indicated by the black vertical line).

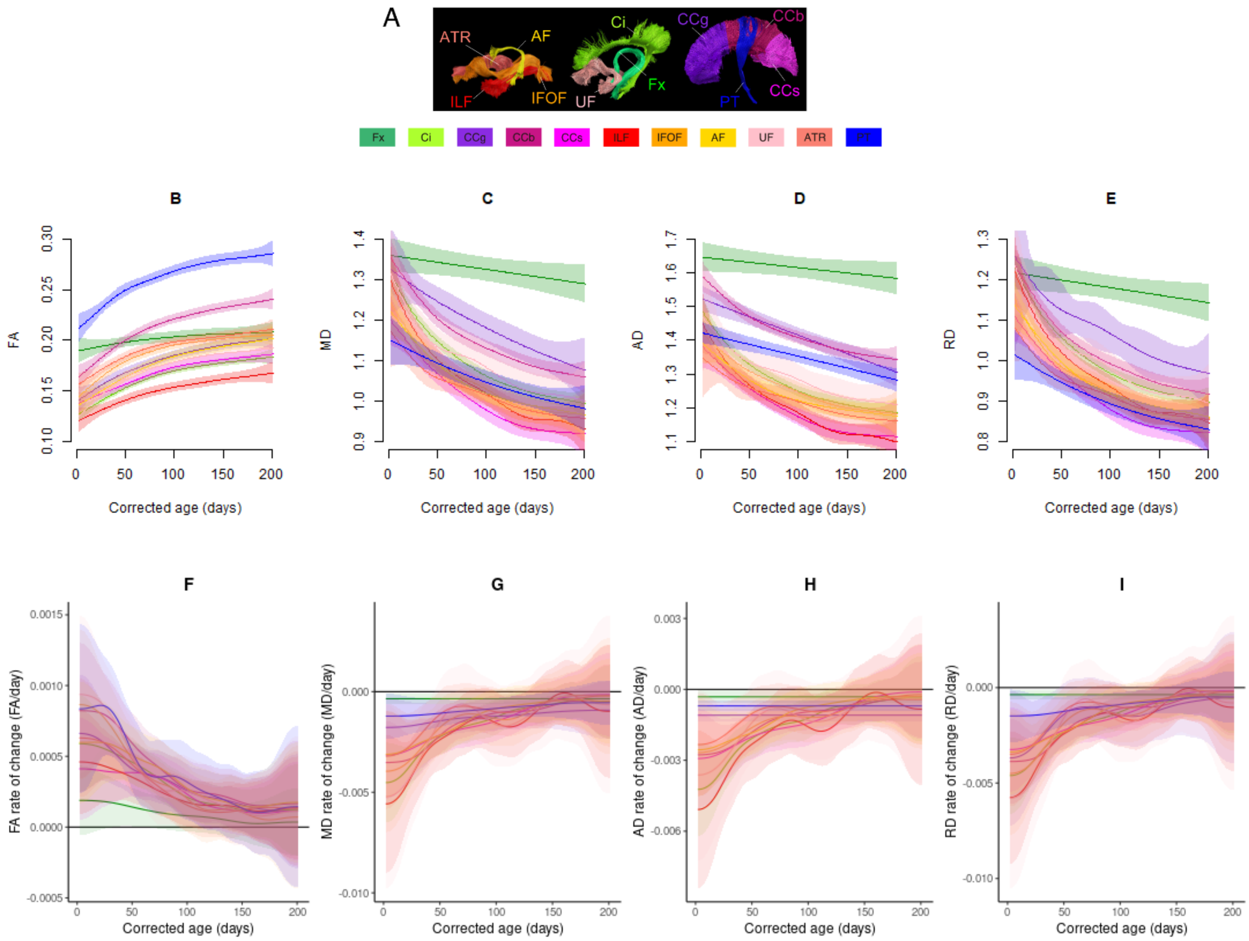

**Supplementary Figure 5. Growth curve (B-E) and growth rates (F-I) of 11 individual WM tracts for four DTI metrics**. Bands represents the 95% simultaneous confidence bands.

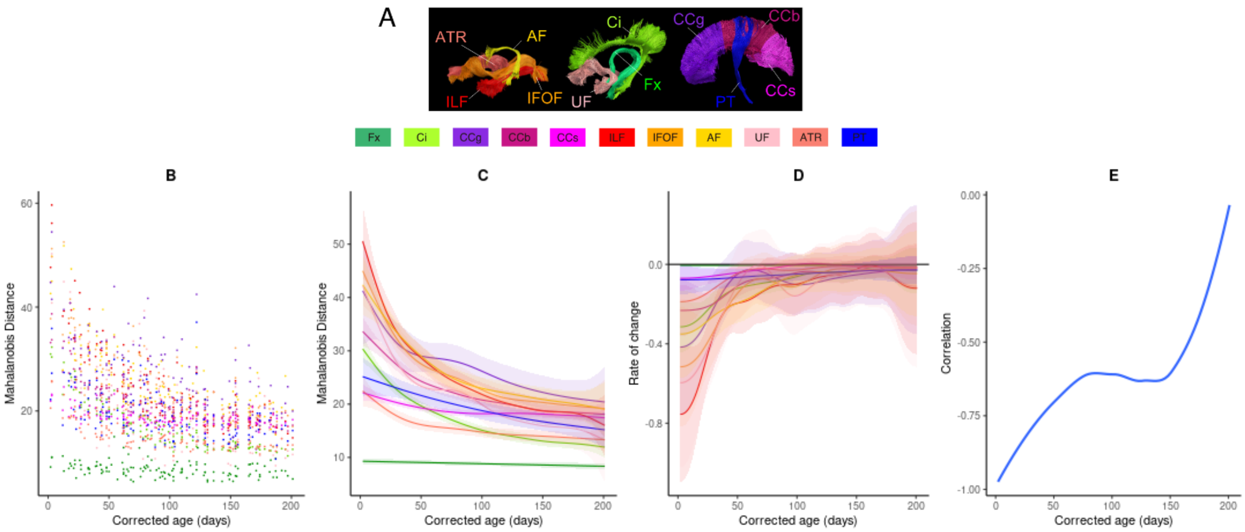


**Supplementary Figure 6. Mahalanobis distance of 11 infant white-matter pathways when compared to adults.** Identical to Fig. 3 in manuscript but with the addition of 95% simultaneous confidence bands.

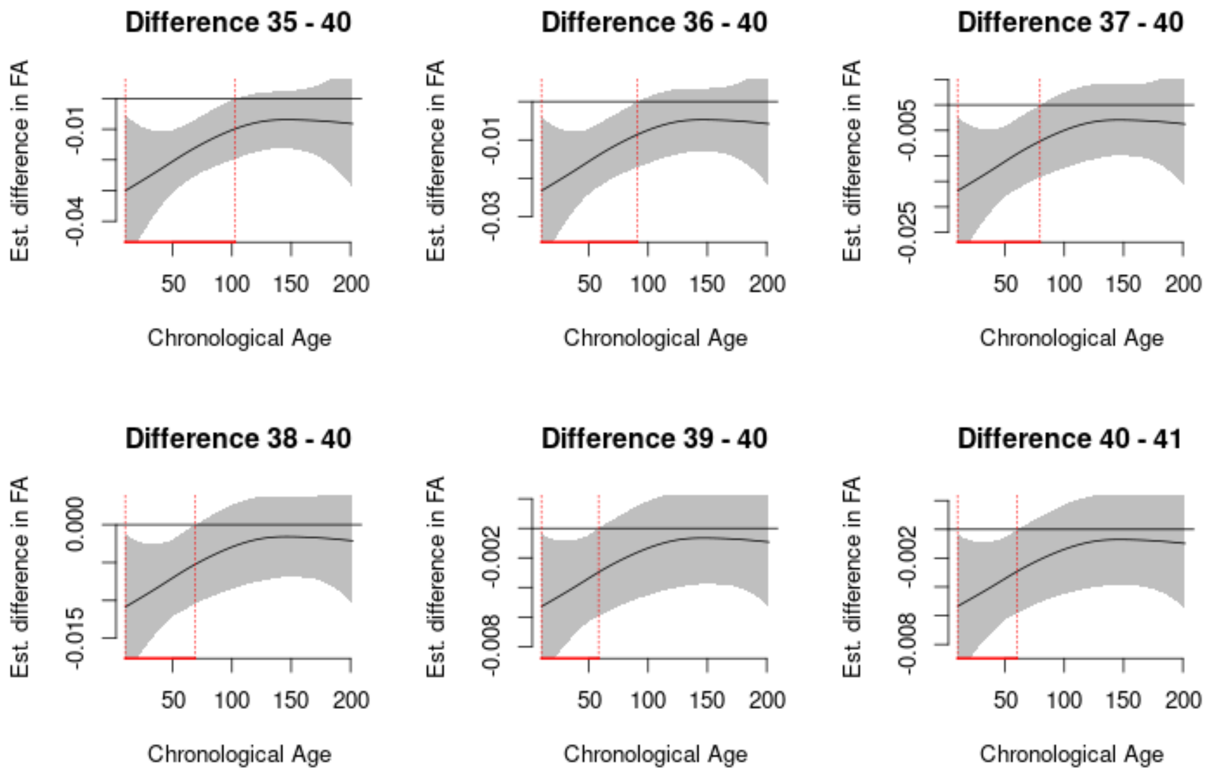


**Supplementary Figure 7. The difference between the chronological age growth curve for gestational age at birth equal to 40 weeks versus gestational age equal to 35, 36, 37, 38, 39, or 41 in whole brain FA.** Red lines on the x-axis denote the windows over which the 95% simultaneous confidence band excludes 0. The curve for GA 41 weeks is significantly higher than GA 40 weeks until 60 days and the curves for GA 39, 38, 37, 36, and 35 weeks are significantly lower than GA 40 weeks until 58, 69, 79, 91, and 103 days, respectively.

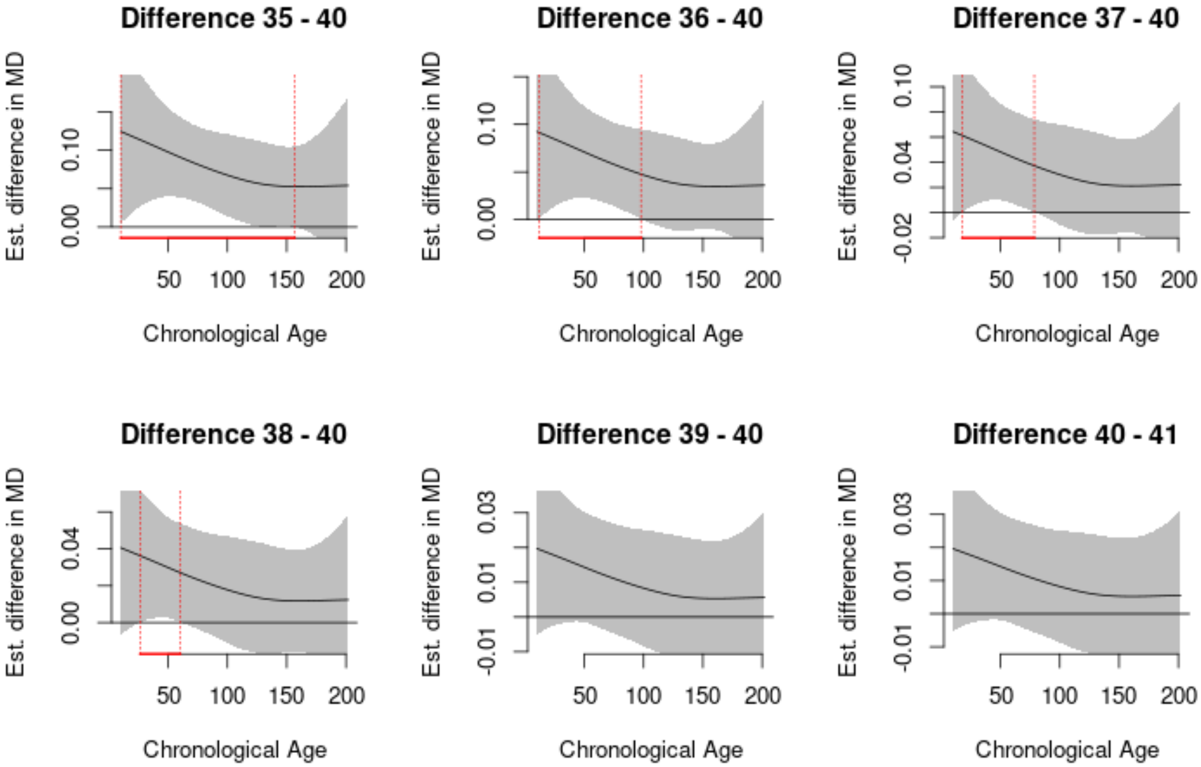


**Supplementary Figure 8. The difference between the chronological age growth curve for gestational age at birth equal to 40 weeks versus gestational age equal to 35, 36, 37, 38, 39, or 41 in whole brain MD.** Red lines on the x-axis denote the windows over which the 95% simultaneous confidence band excludes 0. The curves for GA 38, 37, 36, and 35 weeks are significantly lower than GA 40 weeks until 60, 79, 98, and 157 days, respectively, while the curves for GA 39 and 41 do not significantly differ from GA 40 weeks.

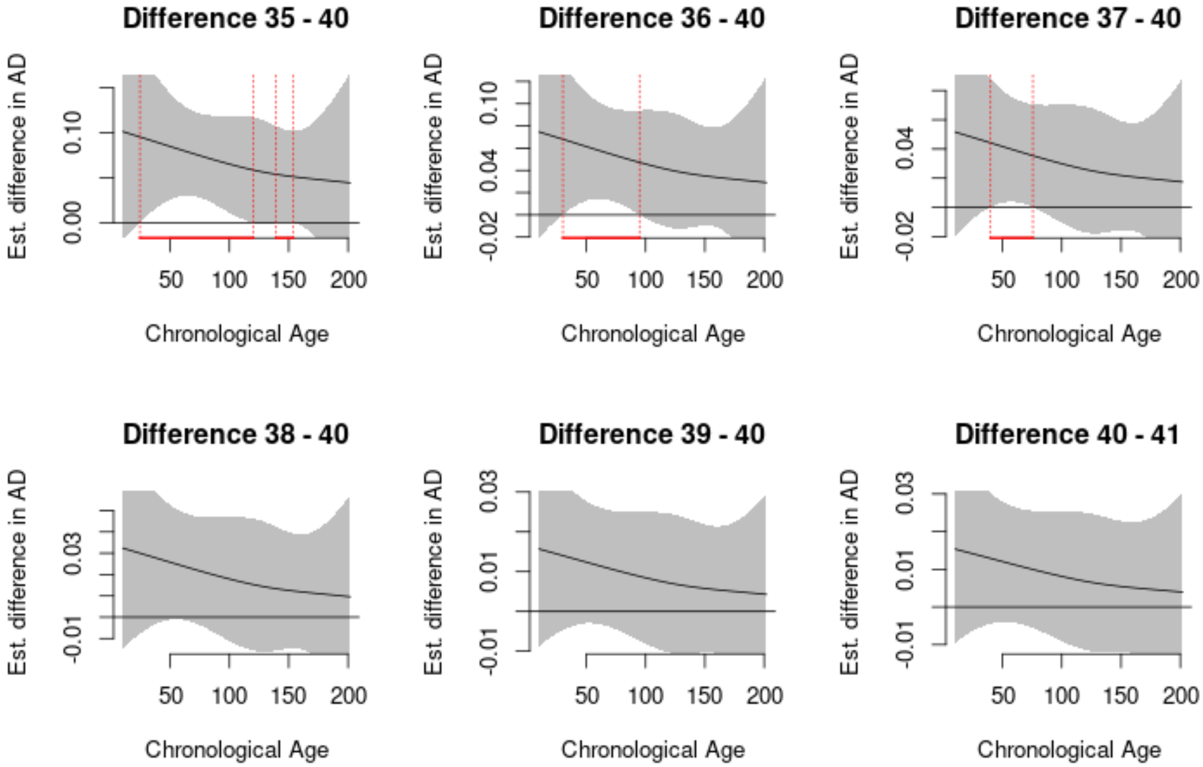


**Supplementary Figure 9. The difference between the chronological age growth curve for gestational age at birth equal to 40 weeks versus gestational age equal to 35, 36, 37, 38, 39, or 41 in whole brain AD.** Red lines on the x-axis denote the windows over which the 95% simultaneous confidence band excludes 0. The curves for GA 37, 36, and 35 weeks have periods during which they significantly differ from GA 40 weeks, with no significant differences after 76, 95, and 154 days, respectively, while the curves for GA 38, 39, and 41 do not significantly differ from GA 40 weeks.

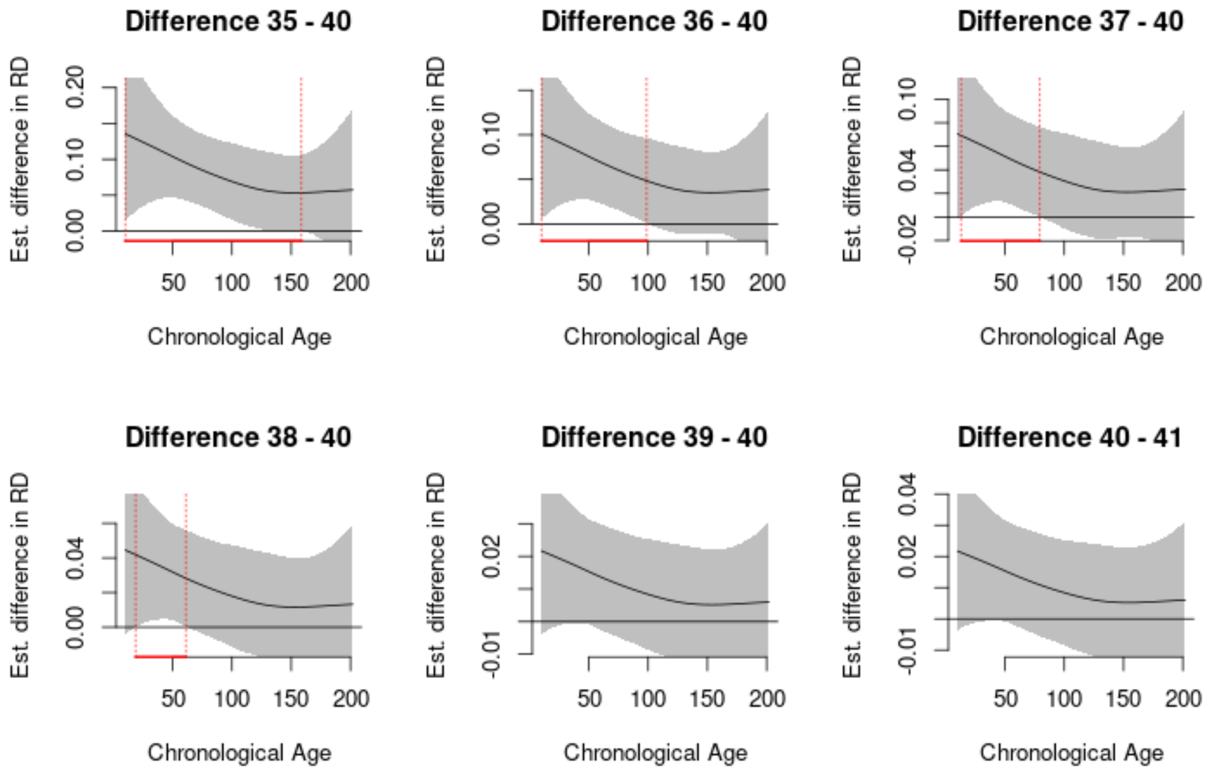


**Supplementary Figure 10. The difference between the chronological age growth curve for gestational age at birth equal to 40 weeks versus gestational age equal to 35, 36, 37, 38, 39, or 41 in whole brain RD.** Red lines on the x-axis denote the windows over which the 95% simultaneous confidence band excludes 0. The curves for GA 38, 37, 36, and 35 weeks are significantly lower than GA 40 weeks until 61, 79, 99, and 159 days, respectively, while the curves for GA 39 and 41 do not significantly differ from GA 40 weeks.

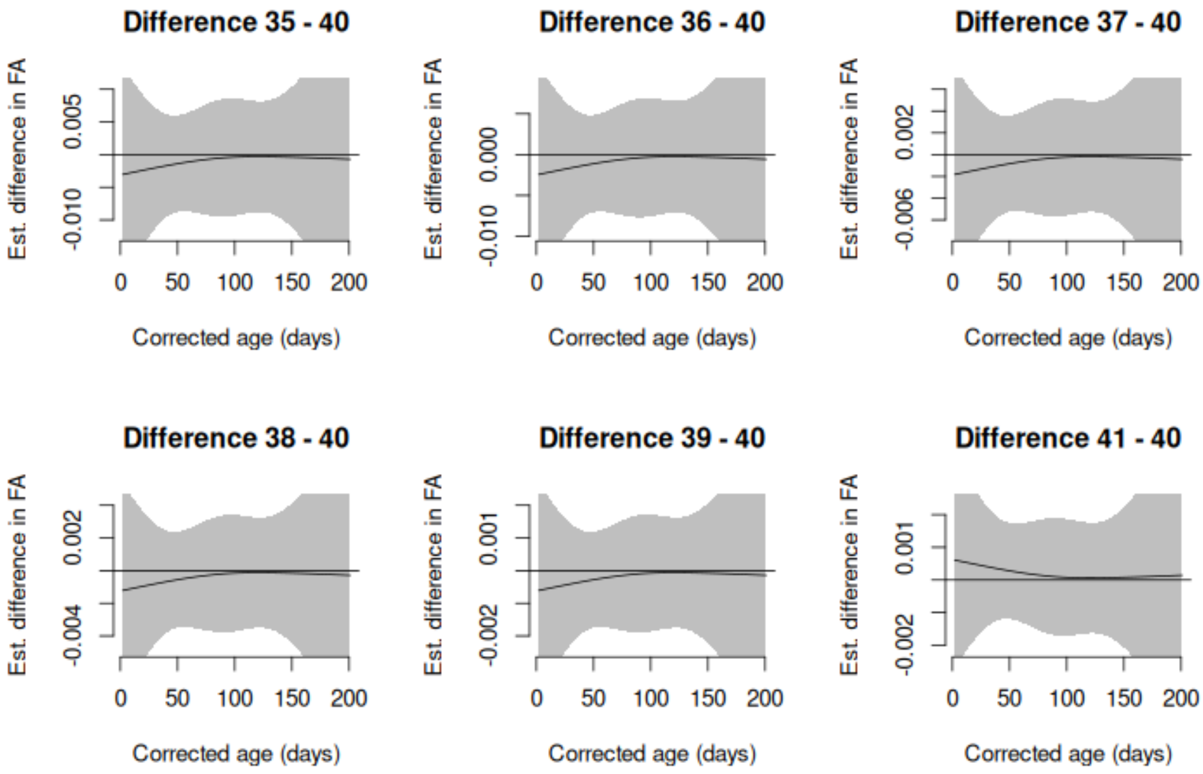


**Supplementary Figure 11. The difference between the corrected-age growth curve for gestational age at birth equal to 40 weeks versus gestational age equal to 35, 36, 37, 38, 39, or 41 in whole brain FA.** Differences are not significant.

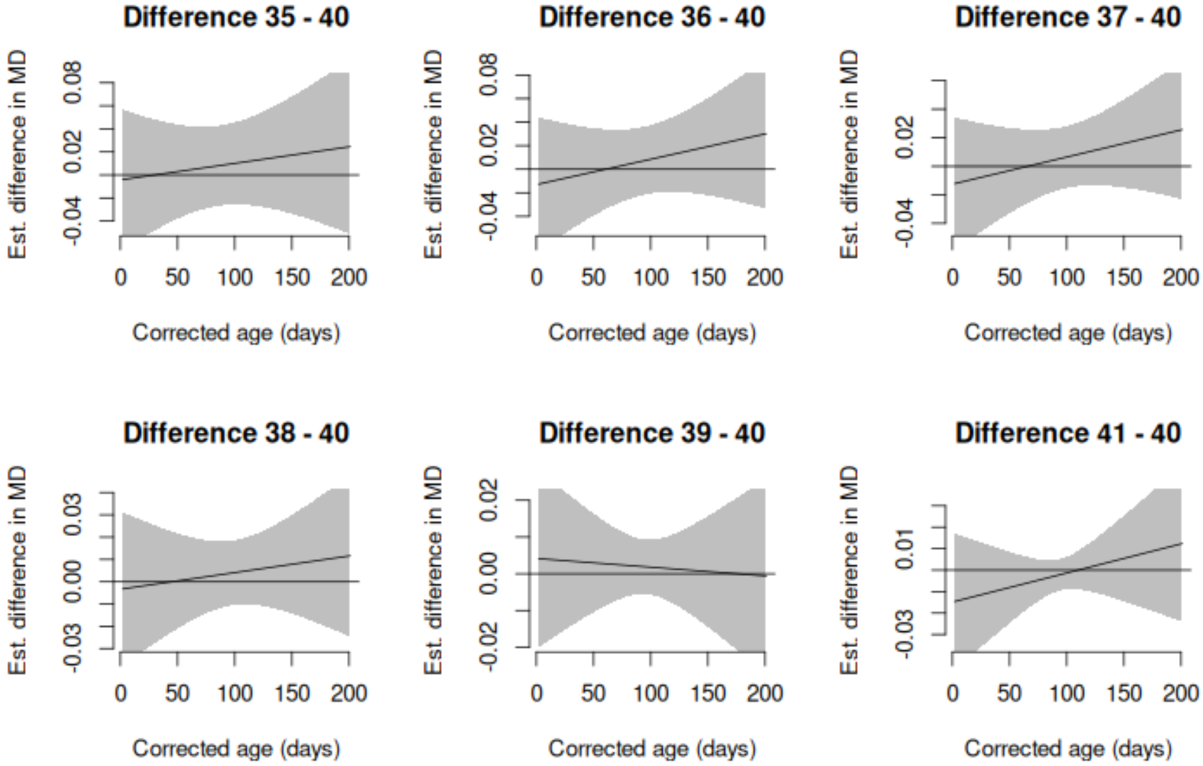


**Supplementary Figure 12. The difference between the corrected-age growth curve for gestational age at birth equal to 40 weeks versus gestational age equal to 35, 36, 37, 38, 39, or 41 in whole brain MD.** Differences are not significant.

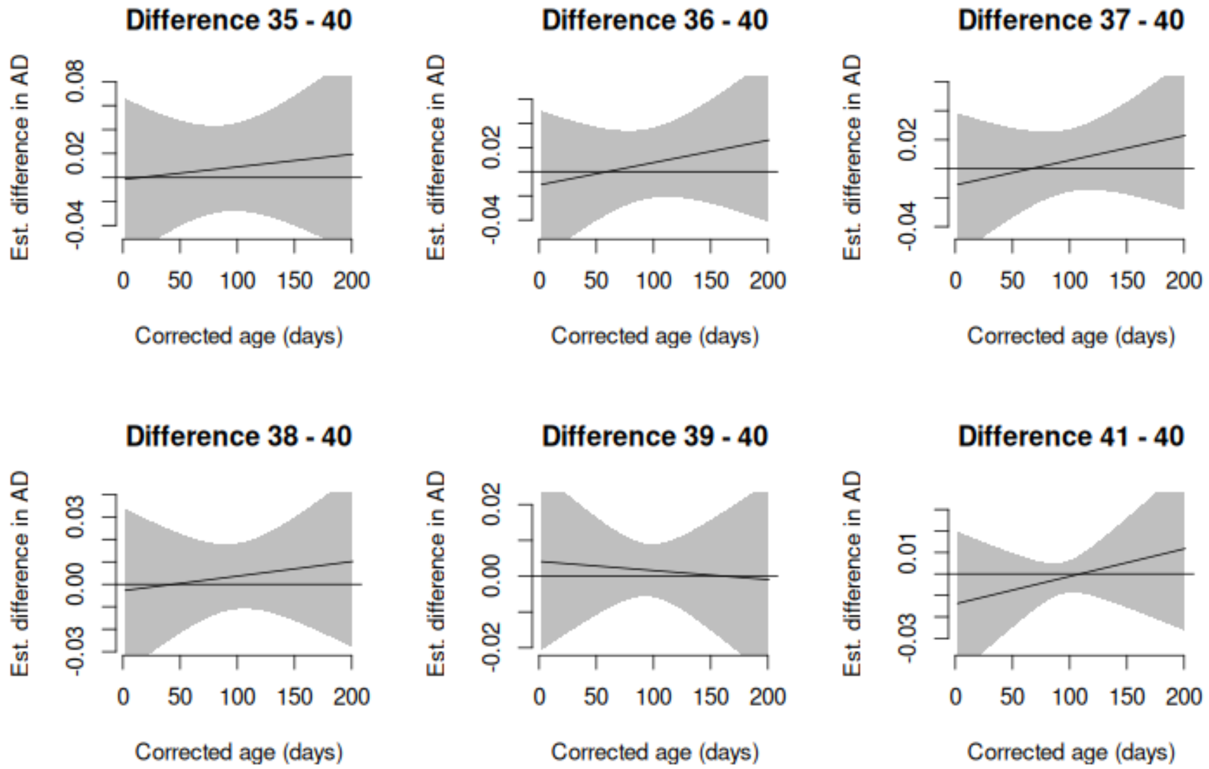


**Supplementary Figure 13. The difference between the corrected-age growth curve for gestational age at birth equal to 40 weeks versus gestational age equal to 35, 36, 37, 38, 39, or 41 in whole brain AD.** Differences are not significant.

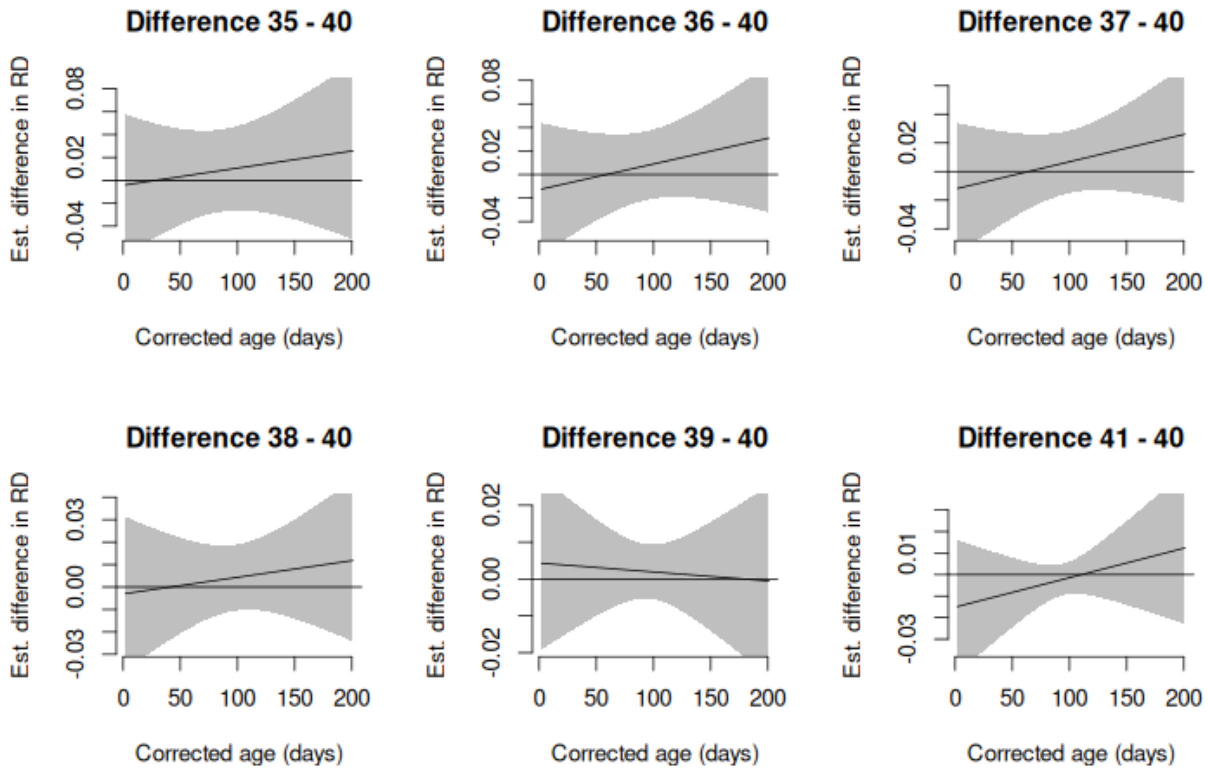


**Supplementary Figure 14. The difference between the corrected-age growth curve for gestational age at birth equal to 40 weeks versus gestational age equal to 35, 36, 37, 38, 39, or 41 in whole brain RD.** Differences are not significant.

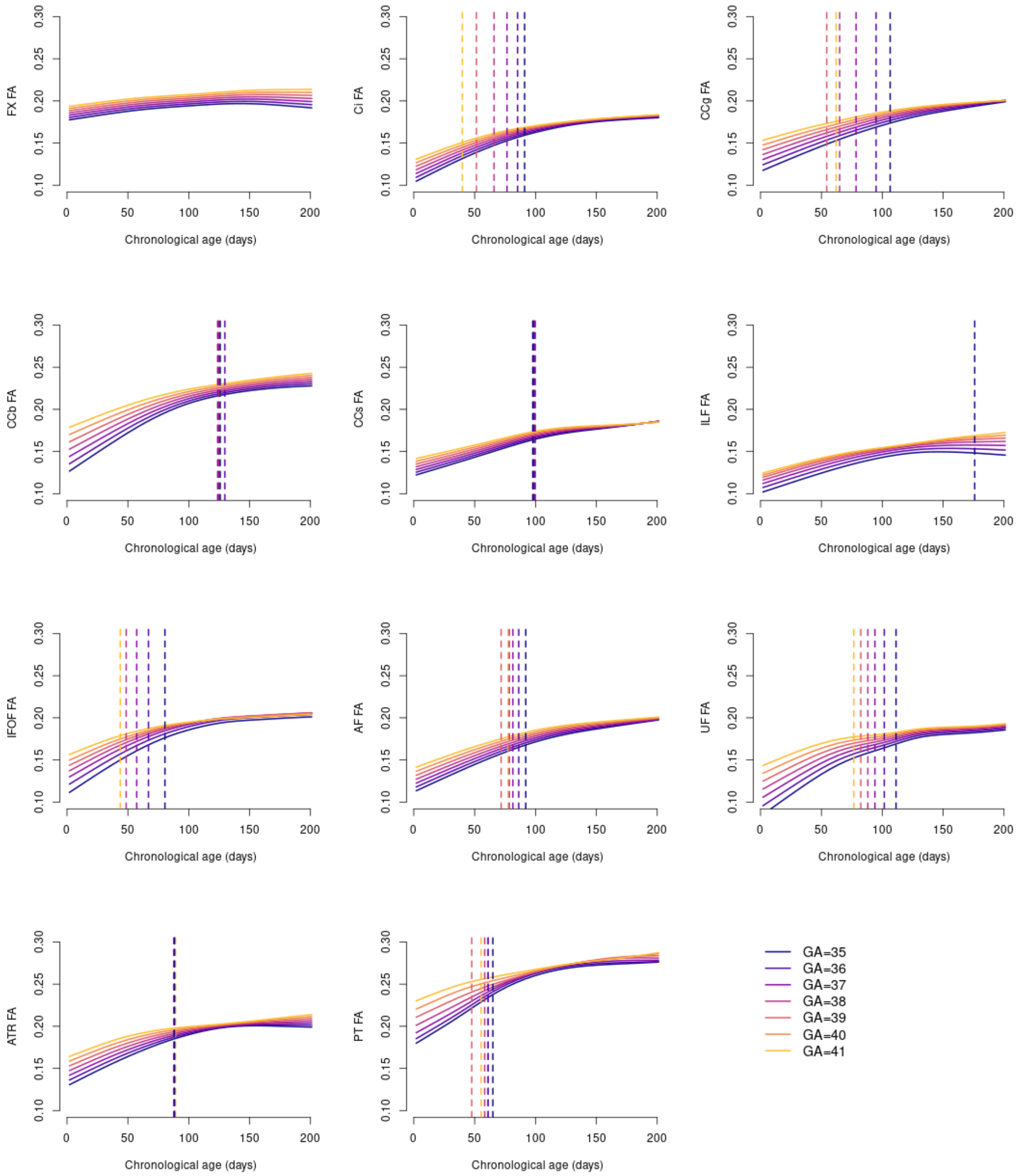


**Supplementary Figure 15.** Effects of gestational age at birth (GA) on infant brain white matter development as indexed by FA. Dashed lines indicate the upper range of the period over which simultaneous confidence bands of the difference curve between the given GA and GA=40 did not include zero. In some instances, simultaneous confidence bands of difference curves contained zero from birth until roughly two weeks, where there were fewer scans (not shown). For example, the simultaneous confidence bands of the PT difference curve did not include zero from 16 to 65 days.

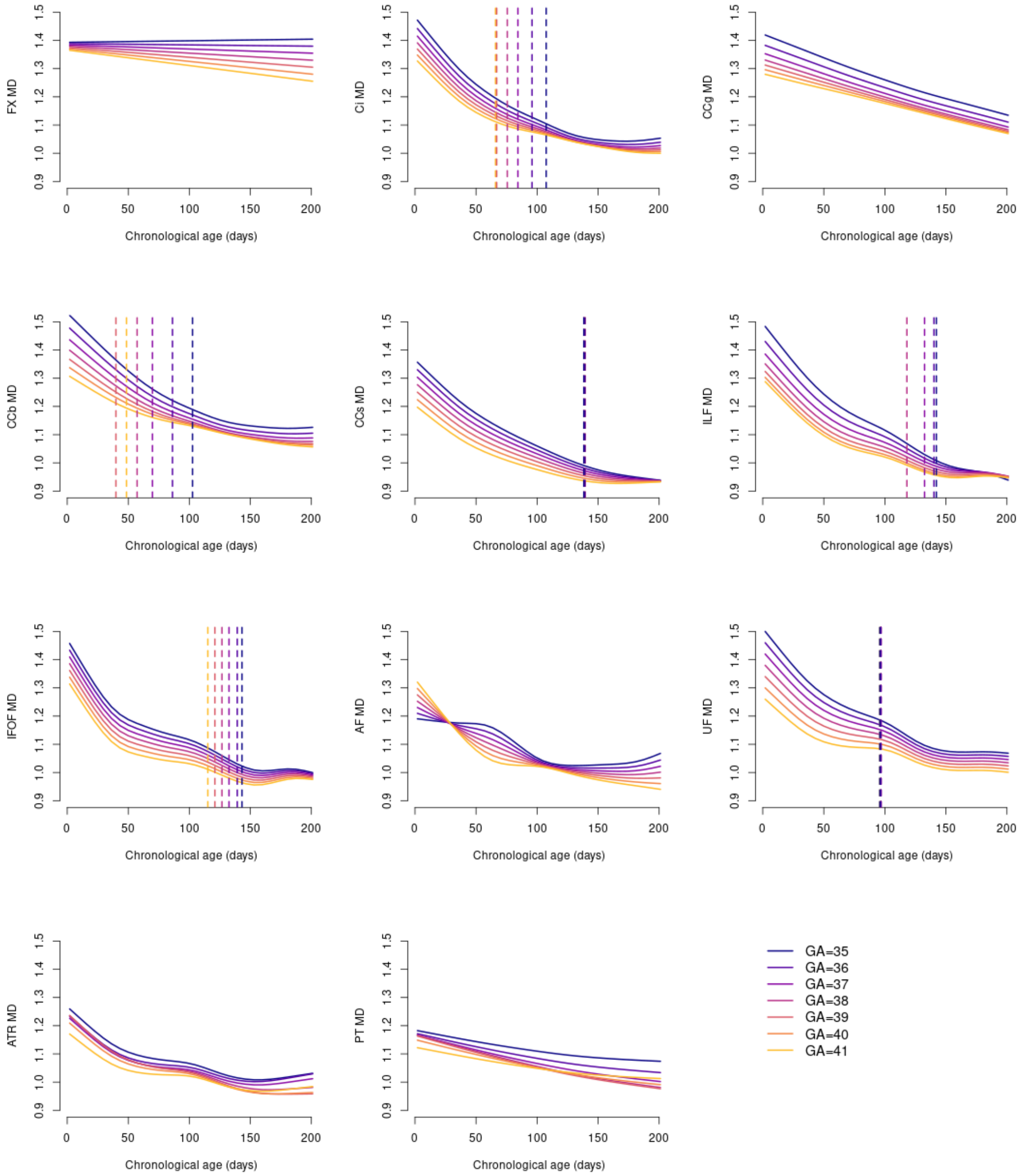


**Supplementary Figure 16.** Effects of gestational age at birth (GA) on infant brain white matter development as indexed by MD. Dashed lines indicate the upper range of the period over which simultaneous confidence bands of the difference curve between the given GA and GA=40 did not include zero.

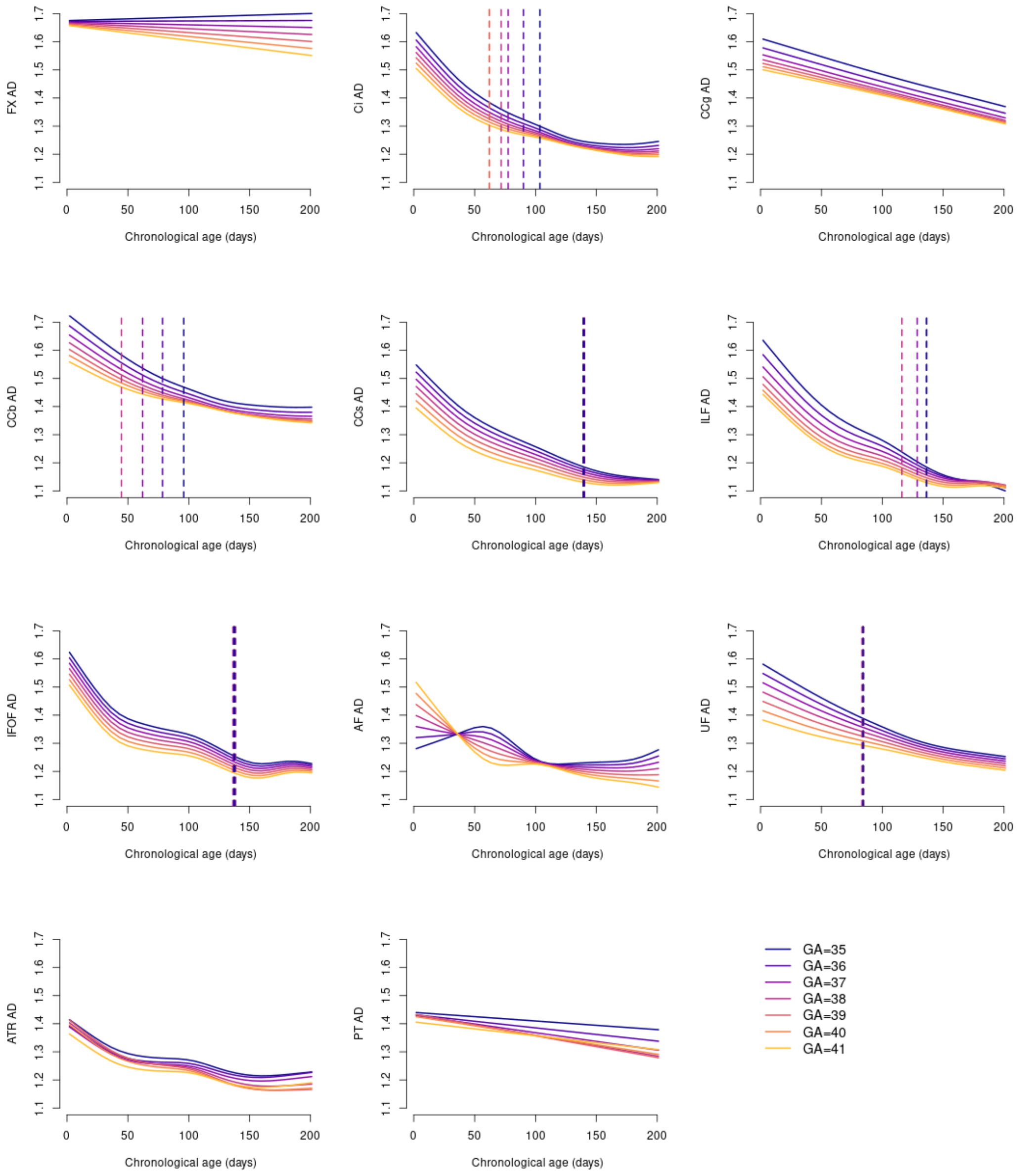


**Supplementary Figure 17.** Effects of gestational age at birth (GA) on infant brain white matter development as indexed by AD. Dashed lines indicate the upper range of the period over which simultaneous confidence bands of the difference curve between the given GA and GA=40 did not include zero.

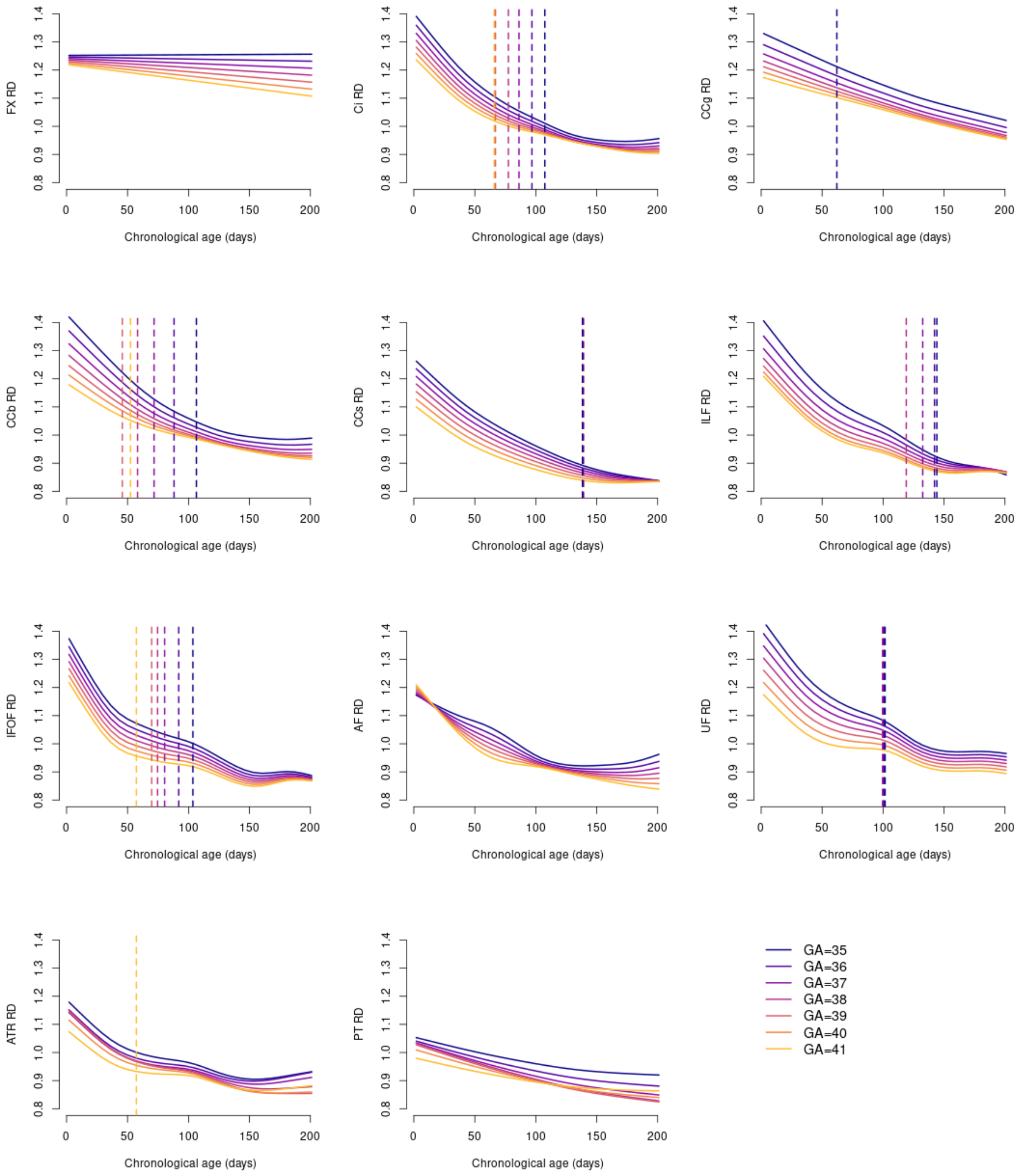


**Supplementary Figure 18.** Effects of gestational age at birth (GA) on infant brain white matter development as indexed by RD. Dashed lines indicate the upper range of the period over which simultaneous confidence bands of the difference curve between the given GA and GA=40 did not include zero.

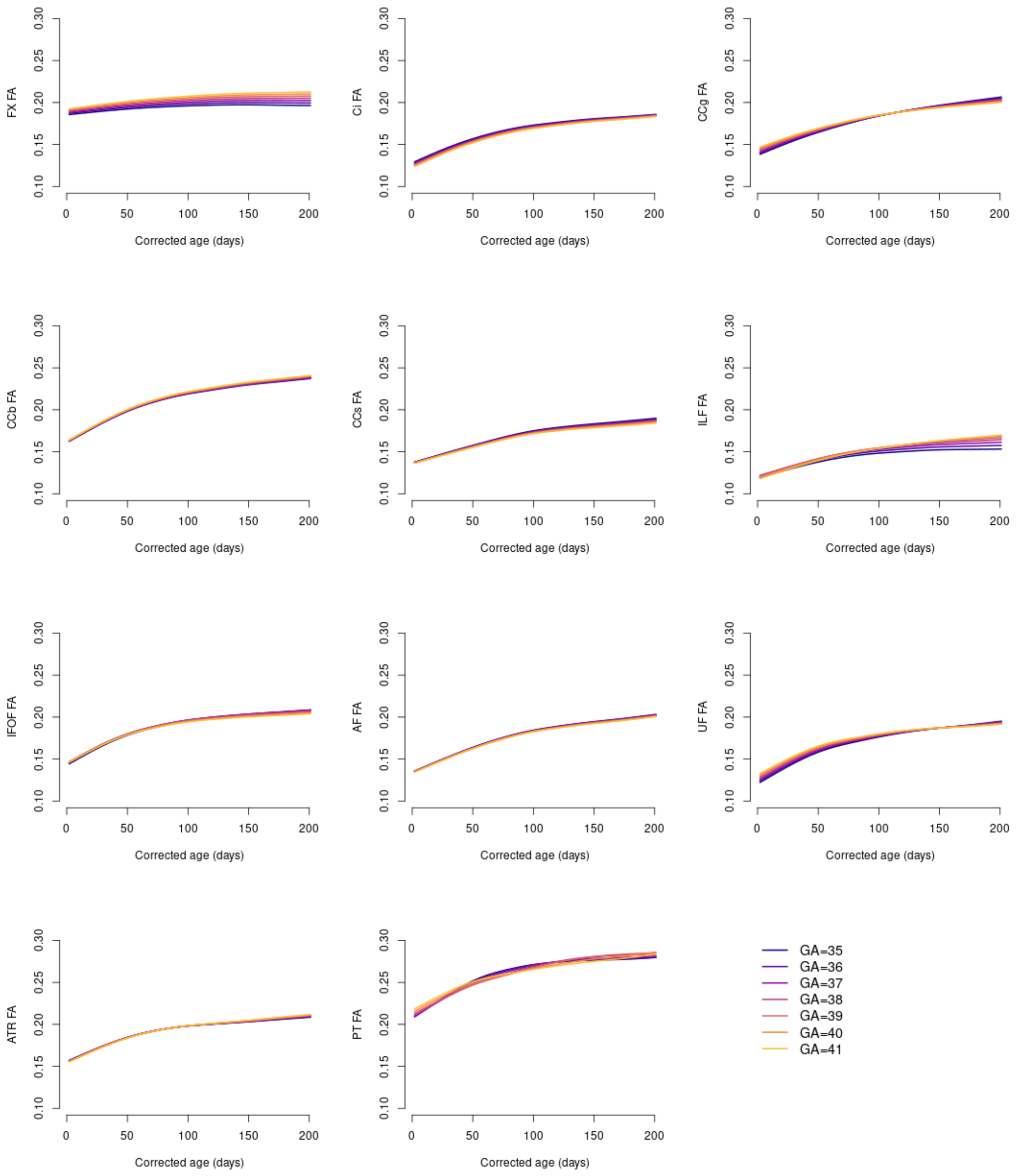


**Supplementary Figure 19. Effects of gestational age at birth (GA) on infant brain white matter development as indexed by FA.** There are no significant differences between age-corrected growth curves for different GA.

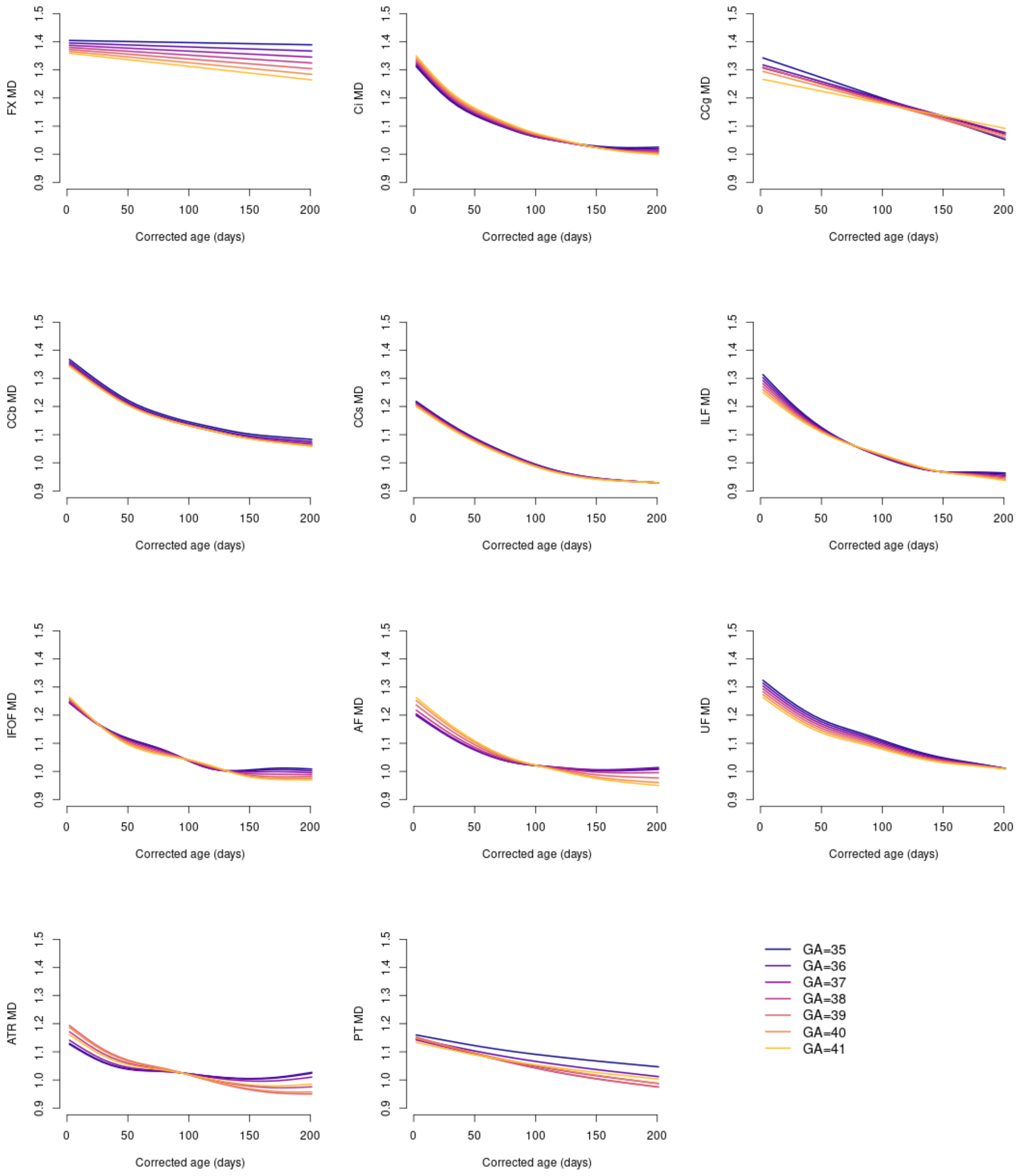


**Supplementary Figure 20. Effects of gestational age at birth (GA) on infant brain white matter development as indexed by MD.** There are no significant differences between age-corrected growth curves for different GA.

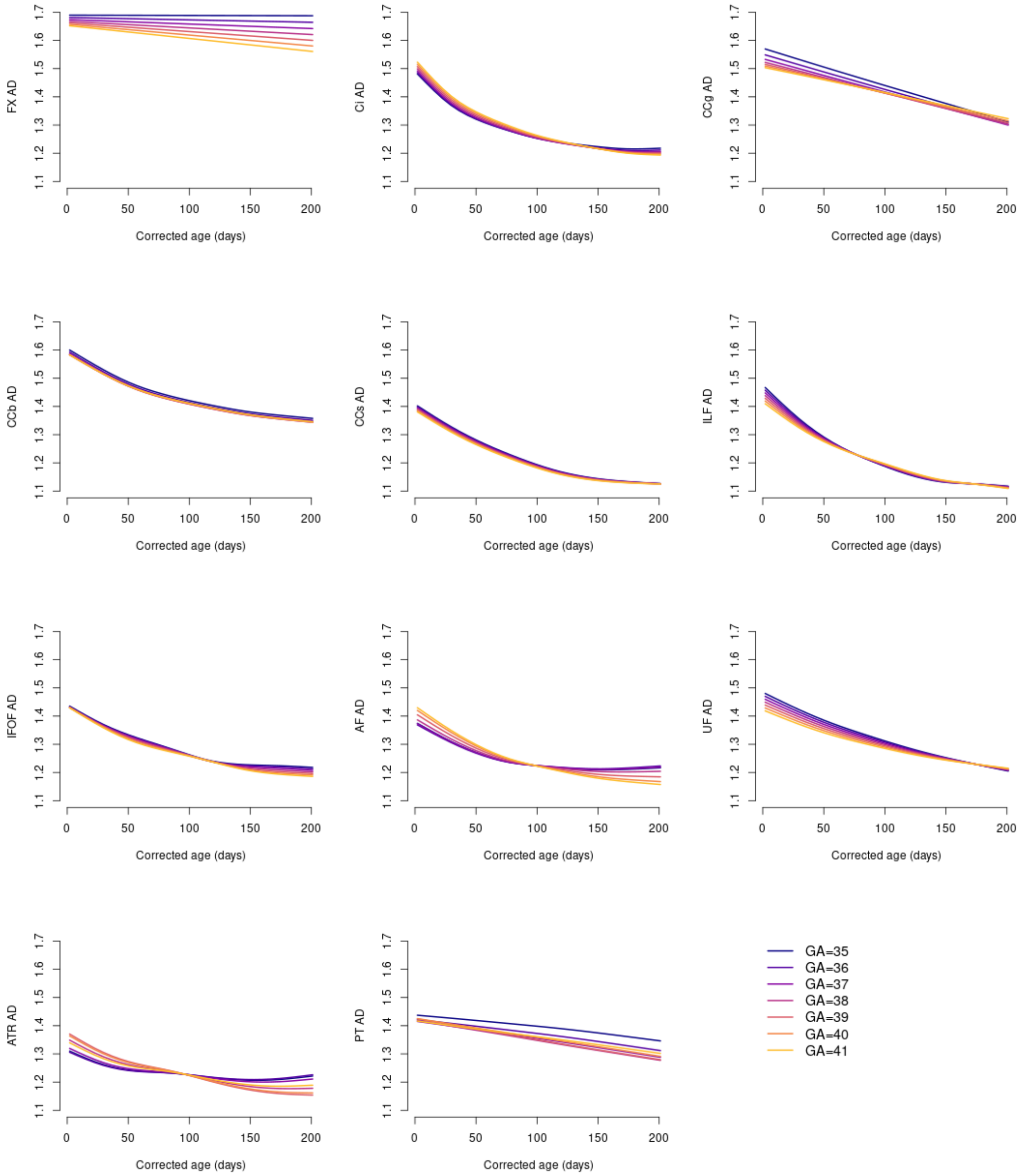


**Supplementary Figure 21. Effects of gestational age at birth (GA) on infant brain white matter development as indexed by AD.** There are no significant differences between age-corrected growth curves for different GA.

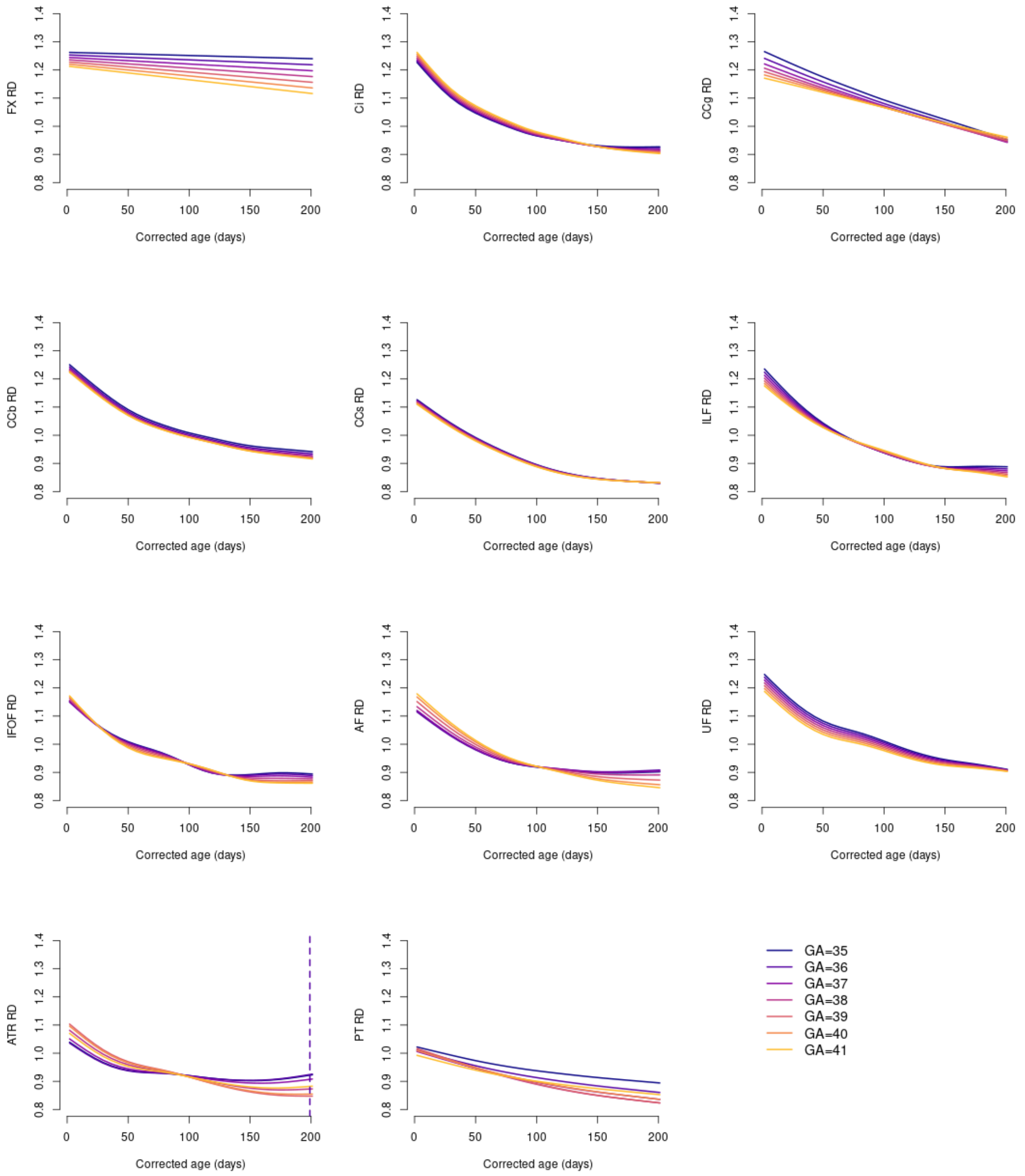


**Supplementary Figure 22. Effects of gestational age at birth (GA) on infant brain white matter development as indexed by RD.** There are few differences between age-corrected growth curves for different GA. The simultaneous confidence band for ATR for the difference in the growth curve for GA=36 and GA=40 is significant from 199 to 201 days.

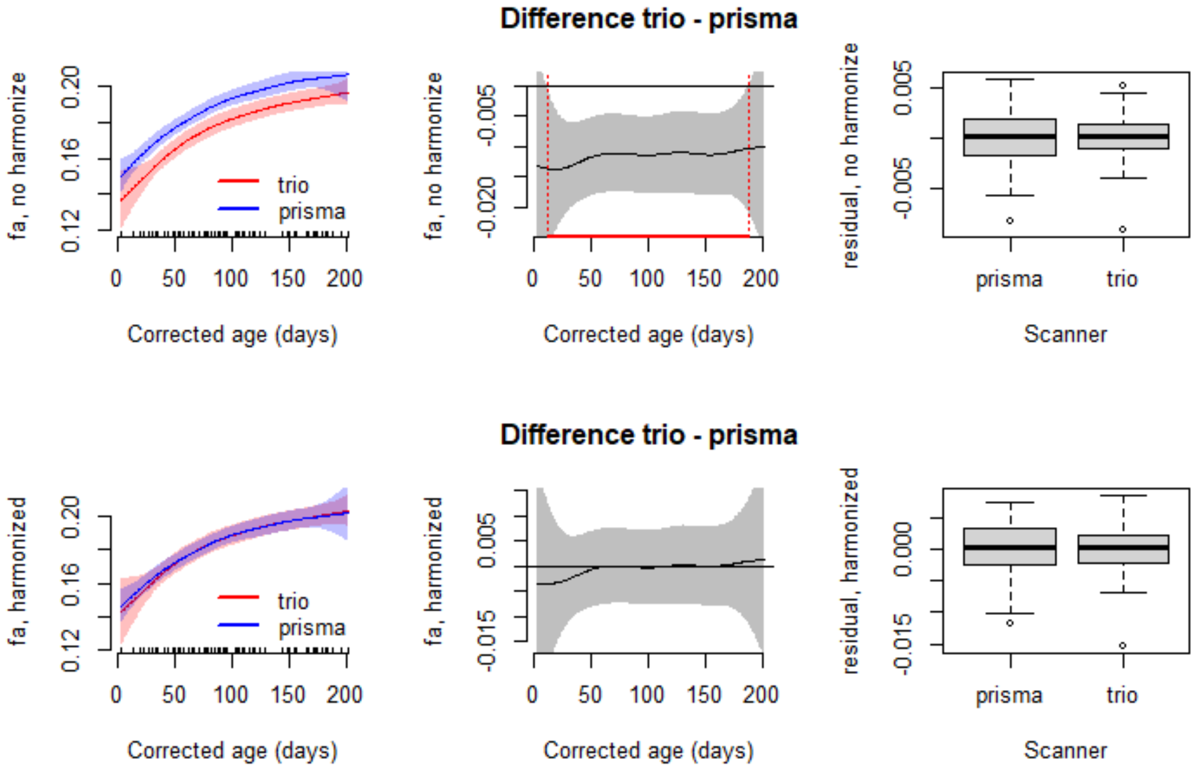


**Supplementary Figure 23. Examining scanner effects and scanner harmonization using longitudinal ComBat in whole-brain FA.** The top row displays the FA growth curves using age corrected for gestational age at birth in the Trio versus PrismaFit, which shows scanner effects. The bottom row shows the curves for each scanner after harmonization.

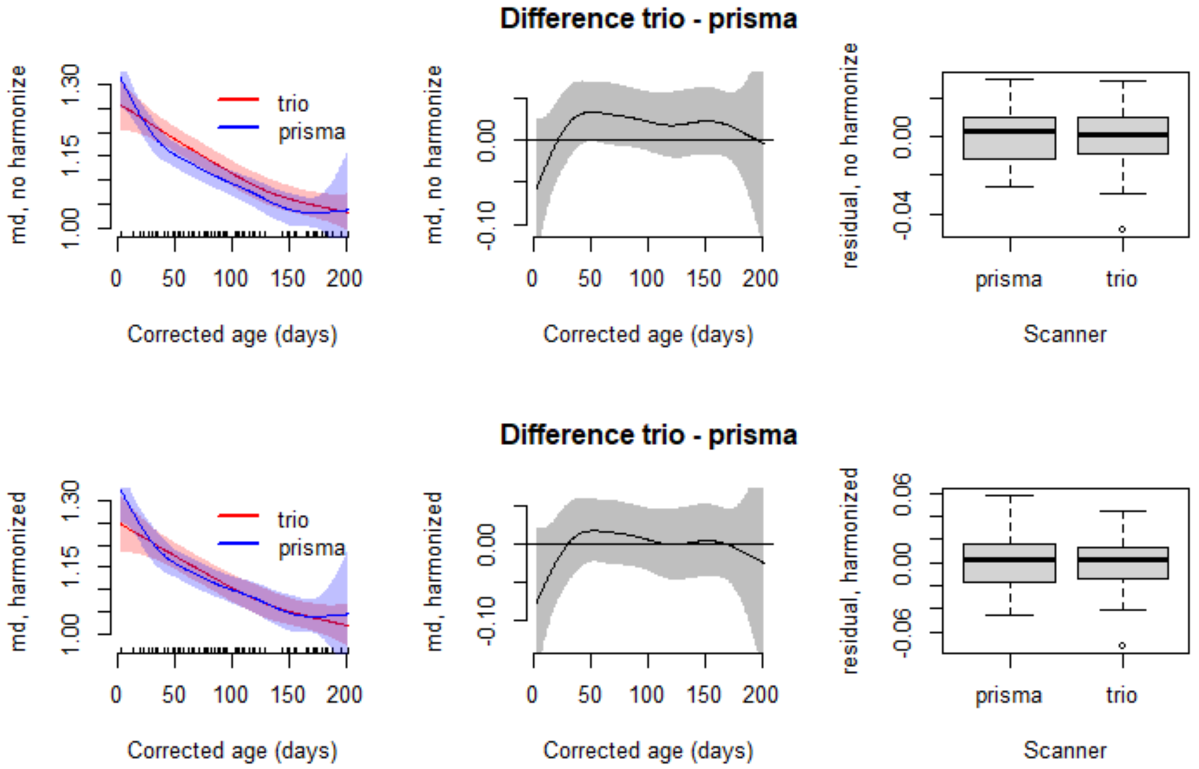


**Supplementary Figure 24. Examining scanner effects and scanner harmonization using longitudinal ComBat in whole-brain MD.** The top row displays the MD growth curves using age corrected for gestational age at birth in the Trio versus PrismaFit. The bottom row shows the curves for each scanner after harmonization.

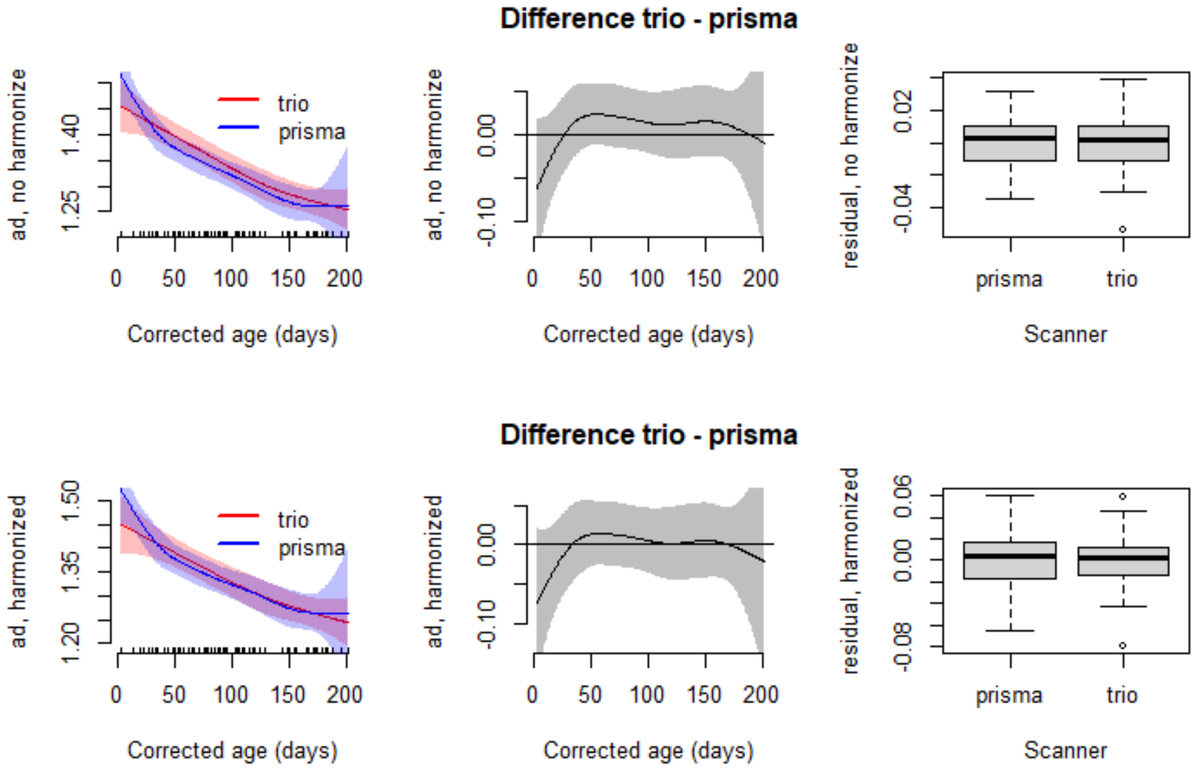


**Supplementary Figure 25. Examining scanner effects and scanner harmonization using longitudinal ComBat in whole-brain AD.** The top row displays the AD growth curves using age corrected for gestational age at birth in the Trio versus PrismaFit. The bottom row shows the curves for each scanner after harmonization.

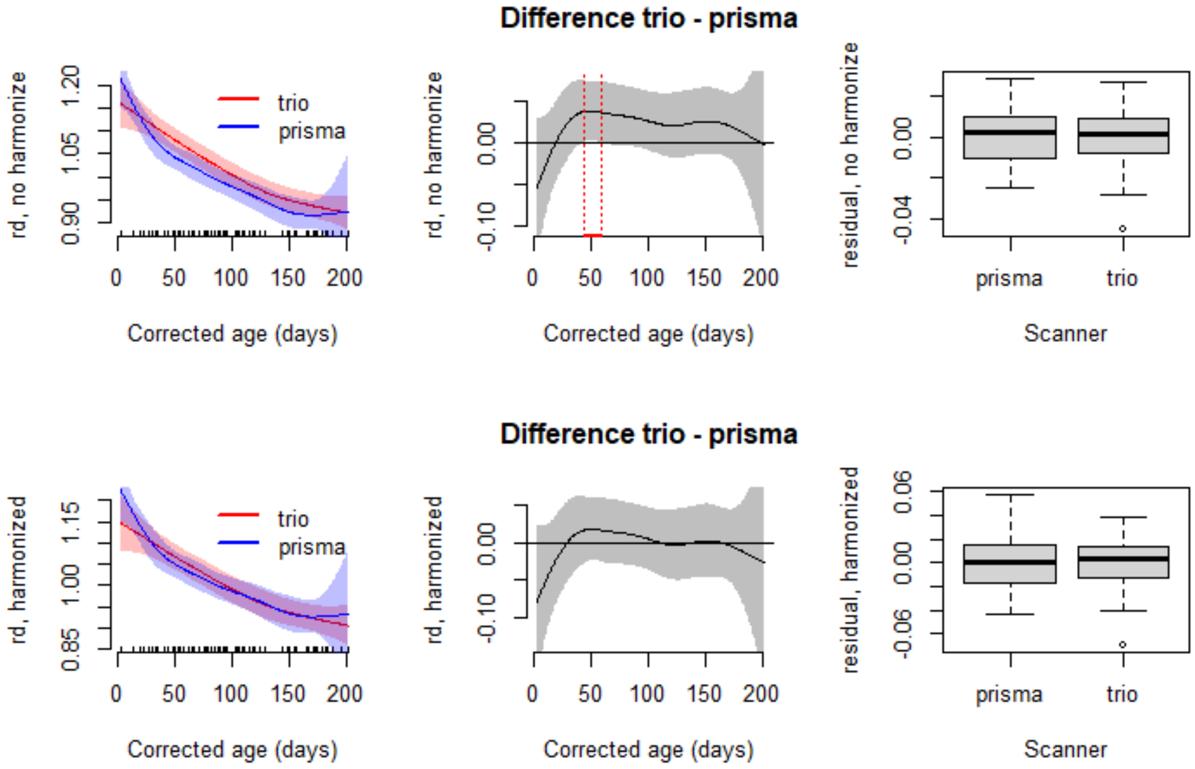


**Supplementary Figure 26. Examining scanner effects and scanner harmonization using longitudinal ComBat in whole-brain RD.** The top row displays the RD growth curves using age corrected for gestational age at birth in the Trio versus PrismaFit. The bottom row shows the curves for each scanner after harmonization.
